# Human Cells for Human Proteins: Isotope Labeling in Mammalian Cells for Functional NMR Studies of Disease-Relevant Proteins

**DOI:** 10.1101/2024.04.09.588766

**Authors:** Philip Rößler, Marco Ruckstuhl, Arnelle Löbbert, Timo Stühlinger, Lucia R. Franchini, Ching-Ju Tsai, Roman Lichtenecker, Binesh Shrestha, Simon H. Rüdisser, Robert Konrat, Gebhard F. X. Schertler, Alvar D. Gossert

## Abstract

In biological and biomedical research the focus progressively moves towards difficult human proteins, which often can only be expressed in higher eukaryotic cells. Nuclear magnetic resonance (NMR) could contribute significantly to the understanding of important proteins as it is one of the most information-rich methods: it allows studying structure, function and dynamics of biomolecules and, importantly, their interactions with natural ligands or drugs. However, to exploit the full potential of NMR, proteins must be isotope labeled. Although expression protocols in e.g. HEK293 cells are often established, isotope labeling is difficult and very expensive. To resolve this disparity, we have developed a comprehensive suite of protocols for isotope labeling in HEK293 cells. We demonstrate uniform 15N and 13C labeling, as well as specific labeling, with special focus on methyl bearing amino acids, including the popular ILV 13C-methyl labeling pattern. Labeling is achieved with a simple laboratory setup and affordable labeling media. These are based on either labeled amino acids, their precursors or amino extracts from microorganisms, like yeast, algae or bacteria.

This enables NMR studies of important, but difficult to produce proteins, like receptors. We therefore expect that these new methods make many highly important proteins accessible to NMR studies and allow exploiting the high information content of this method for accelerating biological and pharmaceutical research.

## Introduction

### Protein production in human cells for NMR studies

To understand human biology and disease, examining human proteins at the molecular level gives insights into their function, their interactions and ultimately can lead to ways of treating a disease using novel therapeutic molecules. Important insights at the molecular level can be obtained with several biophysical techniques [1–7]. All have in common that the target biomolecule must be produced in isolated form. However, many human target proteins are difficult to produce in the preferred laboratory organisms for heterologous expression like *E. coli* bacteria. This is mainly due to the limited folding machinery of *E. coli*, which is not competent to fold more complicated human proteins. This applies mainly to large proteins consisting of several domains, to protein complexes, integral membrane proteins and proteins requiring post-translational modifications. Therefore, for functional and structural studies by cryo-electron microscopy or X-ray diffraction, such proteins are produced in higher eukaryotic cells like insect and mammalian cells. Within the latter, human cells arguably represent the most fitting expression host and for many scientifically and medicinally highly important proteins, expression protocols for human cells are available [8–10]. The fundamental advantages of human cells as expression host are that proteins are correctly folded, assembled into larger complexes and post-translationally modified in a proper way, as they are synthesized and matured by their native production machinery, which has co-evolved with the target proteins.

Nuclear magnetic resonance (NMR) is presently the biophysical technique with the highest information content: molecular interactions, dynamics, function and three-dimensional structure can be studied at the atomic level [11,12]. Importantly, the biomolecules under study do not need to be chemically altered and the molecules can be studied directly in near-physiological solution conditions. The desired level of detail of NMR studies can be gradually adjusted, which reduces the complexity of the experiment and the analysis of NMR data. NMR is therefore highly popular for examining molecular interactions and for functional studies, as little assay development is required and experiments and analysis are comparatively straightforward. However, for resolving signal overlap in large biomolecules, the one requirement NMR has, is that the macromolecule under study is enriched with magnetically active isotopes, typically 13C and 15N. So far, protocols for stable isotope labeling are mainly available for *E. coli* cells and – with some limitations – also for insect cells [13–19]. Therefore, important classes of proteins may be excluded from unique insights that NMR in solution can provide and there is a mismatch between proteins of high interest in biomedical sciences and those amenable to NMR studies. Thus, methods for isotope labeling in human cells are highly sought to enable studying yet inaccessible aspects on highly important proteins.

Protein expression in human cell culture can be carried out in different ways, where the major discerning aspects are: (i) the way of transfection of cells with a gene of interest either transiently or by stable integration into the host’s genome [20], and (ii) the way of culturing cells either adherent on a petri dish or in suspension in a shake flask [21,22]. (i) Human cells in culture allow transient or stable transfection with genes of interest, leading to protocols with either fast turnover or reproducibly high expression levels, respectively. This renders human cells a very attractive system compared to insect cells: Several mutants – e.g. for resonance assignment purposes – can be produced in a matter of 1–2 weeks with transient transfection, while in insect cells the process of virus generation easily requires a month of time. For the central protein in a project, on the other hand, a stable cell line with well-characterized expression characteristics can be kept and expressed whenever needed.

(ii) In order to obtain large quantities of protein, human cells can be adapted to growth in suspension culture, such that large numbers of cells can efficiently be cultivated. The well-established way of culturing HEK293 cells, is growing adherent cells as a monolayer in petri dishes (plates). Culture of adherent cells is relatively inexpensive: media can cost less than 20 EUR/l, and suitable incubators are found in essentially every cell biology laboratory. There are described procedures for isotope labeling with adherent cells [23–29] and a commercial medium for uniform 15N-labeling exists, which is very costly (∼6000 EUR/l). However, the difficult proteins we target here, are typically expressed at levels of 0.2–5 mg/l, such that production of sufficient amounts of protein for an NMR sample, would require growing up to hundred plates with adherent cells grown in this expensive medium.

Since this is very impractical, culturing of cells in suspension is the preferred strategy for production of milligram amounts of protein [30,31]. The major hurdles of suspension culture are however the high costs of media and incubators as well as the lack of proven protocols for isotope labeling in suspension culture. Suspension culture of HEK293 cells requires costly shaker incubators with controlled temperature, humidity and CO2 content, and expensive media (>200 EUR/l) especially designed for suspension culture at relatively high cell densities (>106 cells/ml). With such high starting prices for regular culturing media, custom amino acid-depleted media, as required for isotope labeling, often cost more than 2000 EUR/l, even without counting the cost of isotope labeled amino acids. In summary, isotope labeling in human cells poses many practical and financial obstacles.

We have addressed all these hurdles and have developed a protocol for suspension culture, which is both, enabling and cost-effective at the same time. First, cells are grown in standard cell incubators, just with an added shaking plate. The growth medium, the second cost factor, is replaced by a formulation based on inexpensive media for adherent cells, supplemented with the key ingredients for stable growth in suspension. The final cost of a liter of labeling medium is that of the yeast extract-depleted medium (<200 EUR/l) plus the cost of the isotope labeled amino acids, resulting in costs of 600 EUR/l for uniform 15N labeling, or <250 EUR/l for e.g. methionine 13Cε labeling. Furthermore, we conserve the flexibility offered by mammalian expression systems allowing stable and transient expression.

In this work we developed protocols for obtaining labeling patterns which proved to be the most impactful in previous NMR studies. NMR is mainly used for functional studies by characterizing molecular interactions, conformational changes and/or dynamics[11]. In such studies, either uniformly or selectively 15N-labelled proteins or proteins with 13C-labeled methyl groups of the amino acids Ile, Leu, Val and Met were most often employed. We therefore developed protocols for all these important labeling patterns.

Uniform 15N labeling offers high coverage of a given protein, but spectra tend to get crowded for molecules above 30 kDa. Thus, we also developed amino acid type-selective protocols for e.g. 15N-Val labeling, for reduced crowding of backbone spectra. Amino acid type specific 15N labeling relieves spectral overlap, but often results in limited sensitivity for larger systems. Methyl groups offer superior sensitivity, due to their multiplicity of three and their fast internal motions[32,33]. Thus, 13C-methyl labeling of alanine, using e.g. 2Hα,13C-Ala, yields more sensitive spectra than amide spectra and still confers information on the backbone of a protein. For larger proteins and protein complexes, side chain methyl 13C-labeling is the most suitable labeling pattern. Methyl-bearing amino acids with longer side chains like Thr, Val, Leu, Ile, Met, progressively yield sharper signals due to additional side chain motions, but at the same time typically the coverage of the protein is reduced, with methionine being the least abundant amino acid in this group. However, the sensitivity of methionine methyl spectra is high enough to record methyl signals of particles of 240 kDa without deuteration [34]. We thus implemented 13C-methyl labeling of Ile, Leu, Val and Met.

Depending on the labeling pattern, incorporation of 70–95% of the isotope label is achieved, which is sufficient for 2D spectroscopy, that is used for functional studies. The general strategy is to grow cells in inexpensive non-labelled medium, and then only change to labeling medium for actual protein expression. This approach is economical, but does not allow reaching incorporation above 95%, due to unlabeled metabolites being present in cells. Several passages in the labeling medium would increase the label incorporation asymptotically, but the cost of media rises disproportionally. That is, a second passage in labeling medium might raise incorporation from 75 to 85% but doubles the cost of medium. It therefore makes more sense to produce double the amount of protein with two portions of medium.

Our protocols have been tested with the named labeling patterns, but are surely not limited to these few amino acids. Except for heavily metabolized amino acids like Asn, Gln, Asp and Glu, most amino acids can be labeled with this method. Presently the limitation is mainly the availability of suitably isotope labeled amino acids or amino acid extracts. We thus expect that based on these methods, a large fraction of highly relevant proteins will newly become amenable to NMR studies and thus allow to address pressing questions in biological and medicinal sciences.

## Results

### Cost-effective suspension culture

The step towards suspension culture of mammalian cells represents a significant hurdle, as dedicated shaker-incubators for mammalian cell culture with controlled humidity and CO2 levels are a large investment. Here, we used a basic incubator with active CO2 regulation and passive humidity control by means of a water bath in the base. Such incubators are easy to obtain as second-hand models (Heraeus® and Binder were used here). The shaking function was added by using a shaking plate (Celltron®, Infors, Bottmingen, CH). While maintaining cells and during expression, it is important to monitor cell density and viability. For this task an economic miniaturized microscope with automated cell counting and viability determination software was used (CytoSMART Cell Counter). This entire setup should therefore be affordable for any laboratory.

### Inexpensive media for cell growth in suspension are adaptable for stable isotope labeling

The highly optimized commercial media for suspension culture of HEK293 cells are costly (200–500 EUR/l) and custom amino acid-depleted media are often prohibitively expensive (with quoted prices typically above 2000 EUR/l) or not available at all. Media for suspension culture are inherently more expensive than media for adherent culture, because they must contain more nutrients to support growth to higher cell densities, and they also need to contain factors, which prevent cells from forming aggregates, thus keeping them in suspension – notably an unnatural state for HEK293 cells, without the native, multiple tight contacts between cells.

Instead of using expensive media that are designed for suspension culture, here, an economic medium for adherent culture with low nutrient content (derived from DMEM/Ham’s F-12[35–37]) was used as a base medium and only a few selected ingredients were added to allow cell growth in suspension. To increase the nutrient content, 4 g/l of glucose and 3 g/l of yeast extract were added. In order to maintain cells in suspension a proprietary mixture of lipids was added and albumin at 1 g/l was used to saturate surface receptors on the cells that would either attach to other cells or walls of the culture flask. Such a medium was jointly developed between Novartis AG and Bioconcept AG and resulted in the commercially available medium V3 (V3, Bioconcept AG). When compared to FreeStyle™ 293 medium (Gibco), V3 shows same doubling times of cells (20–25 h), even slightly higher transfection efficiency, but lower maximal cell density (5 vs. 3.5×106 cells/ml). Since protein production is typically initiated at 1×106 cells/ml, the maximally achievable cell density is not so relevant here, and protein yields are essentially the same in the two media. Only in the rare cases where expression proceeds for more than five days, nutrients become limiting. With a price point of <100 EUR/l at the time of writing, this brings the costs for propagating cells to an acceptable level for academic research.

This medium is an ideal platform for isotope labeling, since only the major source of amino acids, the yeast extract, needs to be replaced by an appropriate mixture of amino acids, defined by the desired isotope labeling pattern. Although this labeling medium is a custom medium, the price is affordable (<150 EUR/l, V3-702-I, Bioconcept), since the vendor only needs to exclude one ingredient. The disadvantage is that the medium still contains traces of amino acids from the underlying medium for adherent cell culture (typically tens of milligrams per amino acid, see Fig 2B). Furthermore, 5–10% of fetal bovine serum (FBS) are added to the cell culture, which contain trace amounts of most amino acids. The unlabeled amino acids that are present in low amounts in media are however not so important, since in cell culture there is always a significant carry-over of unlabeled amino acids from the growth culture, which need to be depleted during an incubation period prior to adding the labeled amino acid mixture. By adjusting the duration of the depletion period, also the small amount of unlabeled amino acids are sufficiently reduced, and incorporation levels of 70–95% of the isotope can be achieved. This general approach therefore enables inexpensive isotope labeling with sufficient incorporation and high protein yields.

**Figure 1:**
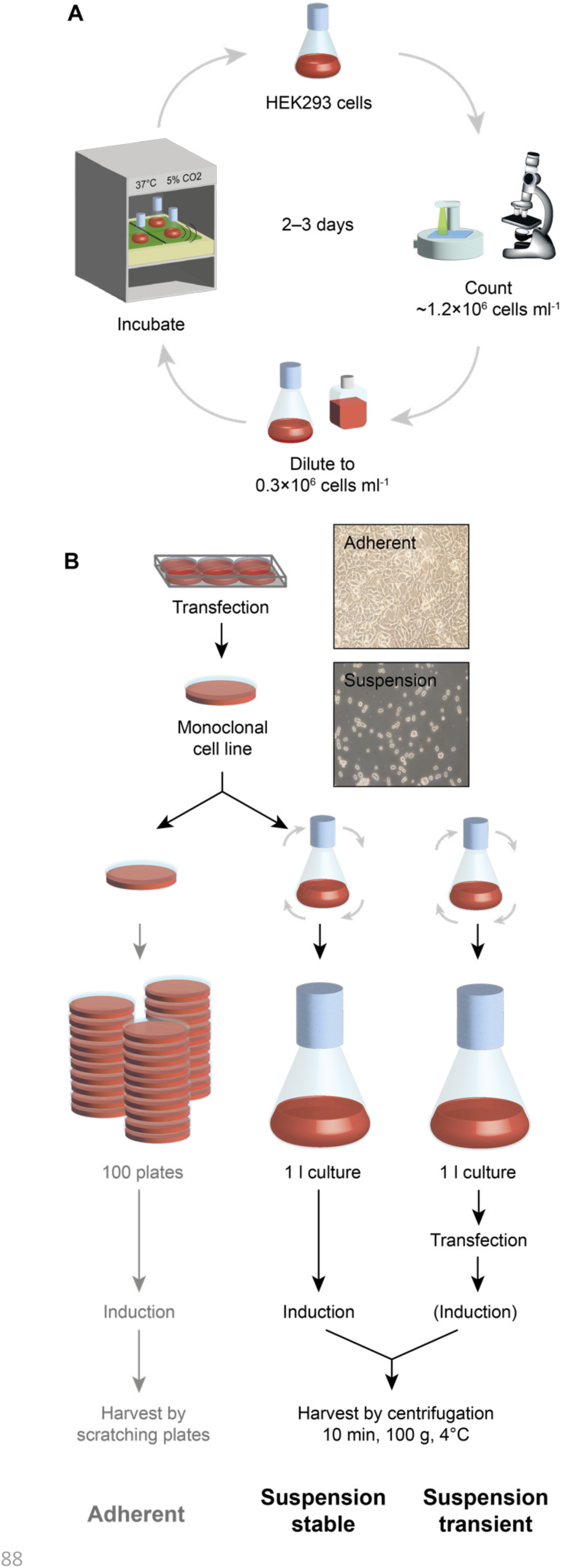
Protein production in HEK293 cells. (A) Maintenance cycle of cells in suspension culture. Cells are kept in a shaker incubator with controlled temperature, humidity and CO2 levels. Every 2–3 days cells are split to 0.3·10^6^ cells/ml. Cell count and viability is monitored using a cell counter and a microscope. (B) Expression of proteins in HEK293 cells. The schematic workflow of protein expression is depicted for adherent cells (left) and stable cell lines in suspension culture (middle) and transiently transfected cells in suspension culture (lower right). After transfection a stably transfected clone is selected by means of antibiotic resistance (2–3 weeks). For expansion, cells are seeded on progressively more plates (adherent culture) or adapted to growth in suspension (suspension culture, 2–3 weeks). Cells can be maintained in suspension for a period of a year, and expanded whenever needed to larger volumes for expression (∼1 week). For transient expression, typically a suspension adapted HEK293T cell line is expanded to the desired volume (∼1 week) and transiently transfected for expression. Depending on the plasmid and the presence of a genomically integrated repressor element in the host cell line, induction may be required. Protein yields tend to be higher in stable cell lines, but transient expression can be accomplished in much shorter time.

**Figure 2:**
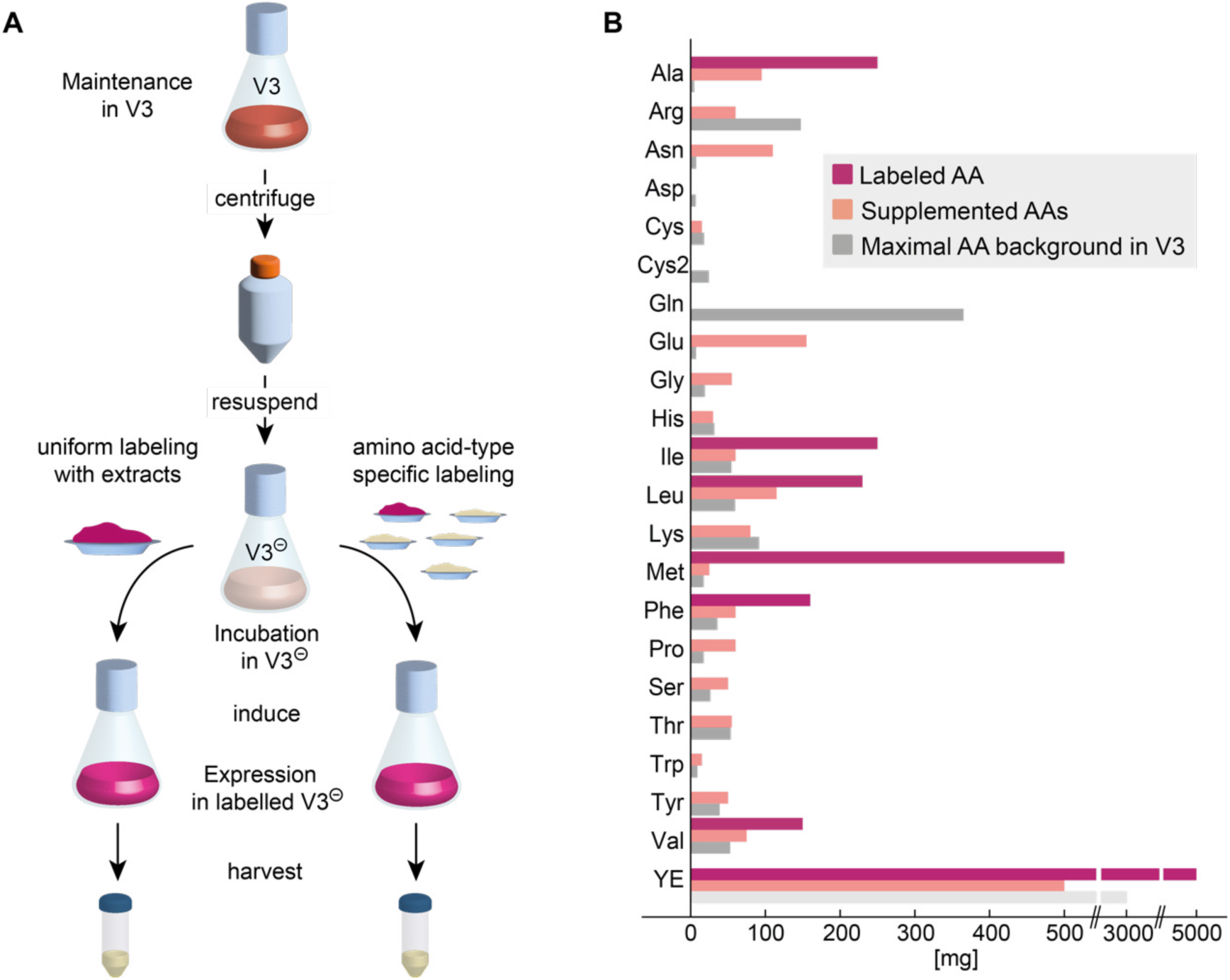
Isotope labeling strategy for HEK293 cells. (A) Cells are expanded in unlabeled V3 medium until the desired volume is reached. Succeeding gentle centrifugation, cells are resuspended in V3 medium without yeast extract (V3⊖). In order to lower the amount of endogenous unlabeled amino acids, cells are incubated for 16 hours in V3⊖. Subsequently, either labeled extract is added for uniform labeling or an amino acid mixture with the appropriate isotope labeling pattern is added to achieve amino acid type specific labeling. This is followed by induction of the protein production for stable cell lines. For transient expression, transfection with plasmid is performed during the incubation period. After 48–60 hours of expression, cells can be harvested. (B) Bar graph displaying the quantity of isotope labeled amino acids and labeled yeast extract (YE) used individually for a desired labeling pattern (dark pink) and unlabeled amino acids and YE (light pink) which are added to V3⊖ to compensate for the lower amino acid content in V3⊖. For comparison the amino acid content of full V3 medium is shown in grey. Note that YE is missing in V3⊖ (light grey).

### Isotope labeling strategies

#### General isotope labeling strategy

The general isotope labeling strategy used here for HEK293 cells is the following: cells are maintained in unlabeled V3 medium and expanded to the desired volume for an expression experiment when needed. When a sufficient amount of culture is available (e.g. 1 l of 106 cells/ml), cells are transferred via gentle centrifugation (maximally 300 g at 37°C) to V3 medium without yeast extract (V3⊖). Here, cells are incubated for 16 hours in order to consume unlabeled amino acids remaining in the cells and in the base medium V3⊖. After this depletion time, an amino acid mixture with the appropriate isotope labeling pattern is added instead of 3 g of yeast extract and protein production is induced. Depending on the desired isotope labeling pattern, the amino acid mixture can be either made up of pure amino acids or a labeled amino acid extract of microbial source, like yeast, algae or bacteria.

#### Uniform 15N labeling

The arguably most popular isotope labeling pattern for NMR studies is uniform 15N- labeling. To this end, V3⊖ medium was supplemented with 5 g of u-15N-labeled yeast extract (autolysates, Cortecnet or Silantes). 5 g were determined to be the best compromise between high incorporation and low price. Except for the slightly higher amino acid content (5 g vs. 3 g of yeast extract), this medium exactly reproduces unlabeled V3 medium, in which the cells were grown, and thus protein yields and growth rates were indistinguishable from non-labeled cultures (see Fig 3B). In Figure 3A a spectrum of uniformly 15N-labeled mEGFP (monomeric enhanced green fluorescent protein [38,39]) produced in this medium is shown. All amino acids were labeled as evident from a comparison with deposited chemical shifts for a similar GFP construct (BMRB entry 5666).

**Figure 3:**
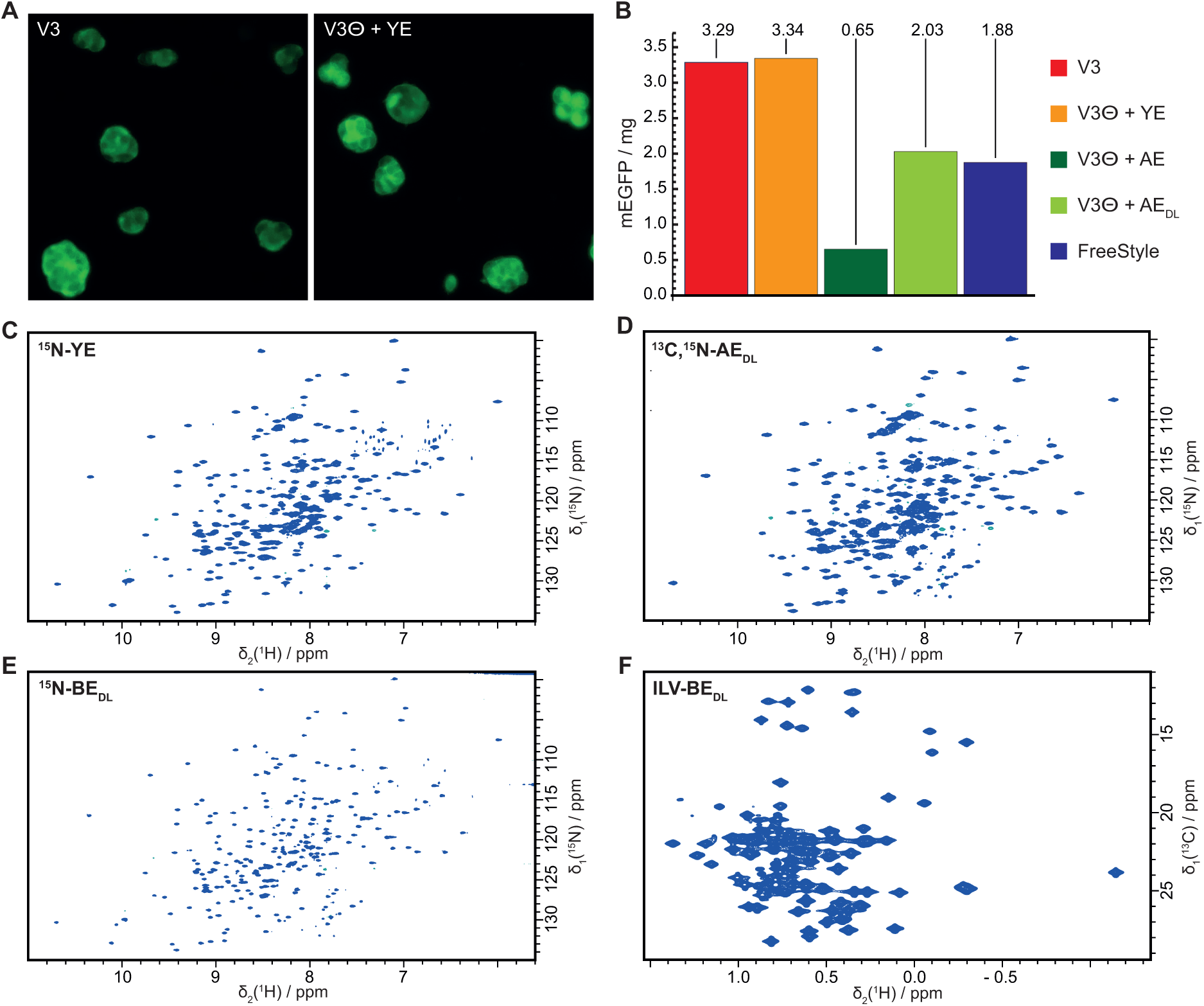
Approaches for uniform isotope labeling in mammalian cells. (A) Microscope images of fluorescent cells from full medium and labeling medium. (B) Yields of mEGFP in mg/l from different media formulations are shown: full V3 medium (V3, red), V3⊖ with yeast extract added back (V3⊖+YE, orange), V3⊖ with algal extract (Celtone) added (V3⊖+AE, dark green), V3⊖ with delipidated (DL) algal extract added (V3⊖+AEDL, bright green) and FreeStyle medium (blue). The yields of mEGFP are in the mg range because it was expressed as a fusion to β1AR and released by protease cleavage. (C–E) 2D TROSY-HSQC spectra of u-15N labeled mEGFP produced in stable HEK293 GnTi- cells either with (C) 15N labeled yeast extract or (D) delipidated 13C,15N-labeled algal extract (ISOGRO, Sigma) All amino acid types were labeled, since identical signals are visible as reported from *E. coli* expression. Incorporation was 70%. (E and F) 15N and 13C HSQCs of ILV-labeled mEGFP produced with *E. coli* extracts with the same labeling pattern.

This approach is very economic, since the isotope costs only amount to 5 g of labeled yeast extract. When compared to the only commercially available medium (BIOEXPRESS- 6000, CIL) with a price point > 6000 EUR/l, the savings are more than ten-fold. Further, the commercial medium is only available for adherent culture, such that there are additional savings in workload, as one liter of easy-to-handle suspension culture corresponds to about hundred plates of adherent culture.

An alternative source of isotope labeled amino acids are algal extracts, which presently are commercially available with more diverse isotope labeling patterns than yeast extracts. We tested the products ISOGRO (Merck) and Celtone (CIL). Since algal extracts have a higher amino acid content than yeast extract (∼60% vs. ∼35%, respectively), only 4 g/l of algal extracts were employed. However, algal extracts severely impaired cell growth and resulted in nearly 90% reduced protein yields. Yields could be restored to about 50%–60% by de-lipidation of the extract (Figure 3C, SI Protocol 4) [29]. De- lipidated algal extracts are therefore an alternative to yeast extracts, enabling more economic access to 2H and 13C isotope labeling.

Up to this point, only commercially available extracts of isotope labeled amino acids were considered, due to the ease of accessibility and the relatively little work involved in the laboratory to prepare such media [40–42]. Preparing isotope labeled amino acid extracts on site is laborious, but potentially allows accessing alternative labeling patterns, for example ILV-labeling [24,43]. To this end, we grew *E. coli* cells in uniform 15N-labeled form with selective methyl 13C-labeling of Ile, Leu and Val methyl groups. In contrast to the original ILV-labeling protocol[44], the precursors α-keto-isovalerate and α-keto- butyrate were added from the beginning of the culture and at higher concentrations (180 and 160 mg/l, respectively), in order to fully label all *E. coli* cell mass. After digestion of the lysed cells with different proteases (Figure 3E and F, SI Protocol 3) and performing the same de-lipidation protocol as for algal extracts, we were able to employ this *E. coli* extract in mammalian growth media. The GFP protein produced in this way in mammalian cells, faithfully reproduced the labeling pattern form *E. coli*, albeit with lower incorporation (∼66%), such that with this protocol we were capable of producing ILV- 13C-methyl labeling in mammalian cells.

In summary, several alternatives for economic amino acid mixtures are there, which can be employed depending on the desired isotope labeling pattern. For uniform 15N labeling yeast extract is preferred because nearly two-fold higher yields can presently be obtained, than with other extracts. For uniform 13C labeling algal extracts are fundamentally more economic, since algae can grow on 13CO2 and don’t require pricy 13C- labeled glucose, and, finally, for more advanced labeling patterns *E. coli* is currently the best option.

#### Amino acid-type specific labeling by replacement

With uniform labeling full coverage of a protein can be obtained, however, especially for larger proteins, spectra tend to become very crowded. In such cases, only labeling one or two amino acid types is a more promising alternative. The generalized approach for amino acid type specific labeling, is using V3⊖ medium and replacing the yeast extract with an amino acid mixture reflecting the desired isotope labeling pattern. Instead of yeast extract, the isotope labeled amino acid(s) and a mixture of the remaining amino acids is added, each at the corresponding amount present in 3 g of yeast extract (SI Table 1 and Fig 2B). The amount of the isotope labeled amino acid is adjusted such, that it has a final concentration of at least 3-fold the concentration of the unlabeled amino acid in the base medium before starvation.

This approach is exemplified with 15N-valine labeling of mEGFP, where 150 mg/l of 15N- Val was employed in the context of V3⊖ medium supplemented with the mixture of unlabeled amino acids shown in the SI and figure 2B. The resulting simplified spectrum is shown in Figure 4, where all valine resonances are visible. Interestingly, a small fraction of isotope scrambling is visible, indicating residual metabolic acitivty of mammalian cells. This indicates that potentially isotope labeling starting from simpler precursors may also be possible in these higher cells. This avenue was explored further below.

**Figure 4:**
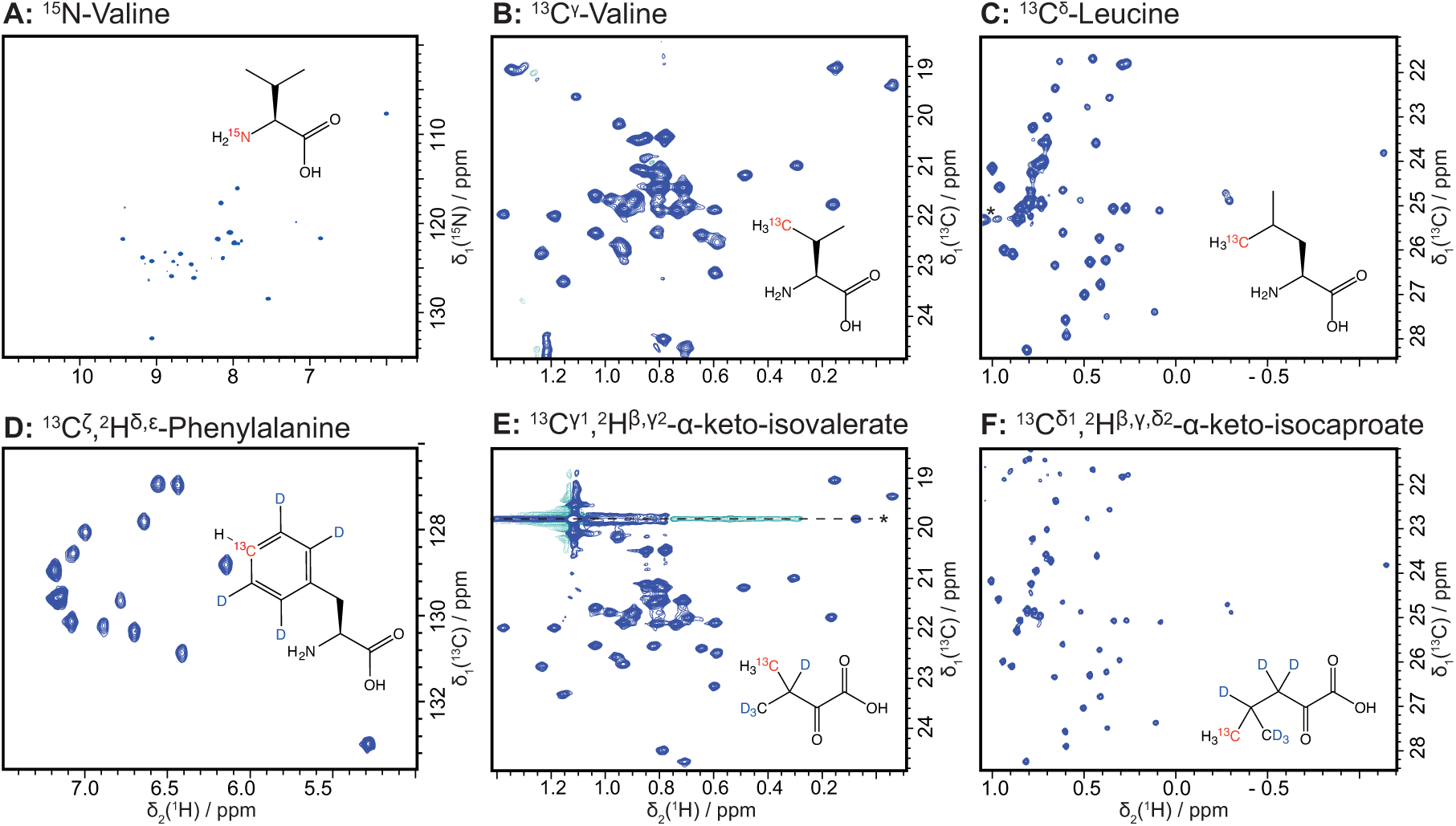
Amino acid type-specific labeling using the 15N, 13C, and 2H isotope labeled reagents indicated on the right in each spectrum. 2D [15N,1H] and [13C,1H] correlation spectra of mEGFP produced in stably transfected HEK293 GnTi- cells are shown. Proteins were produced using either the full amino acid (A–D) or α-keto-acid precursors (E and F). See text and SI for exact experimental conditions.

For larger proteins however, [13C,1H] correlation spectra of methyl groups have proven to be the most sensitive approach. Therefore, 13C-labeling of methyl groups of Ile, Leu, Val and Met is highly important in this context. Here we demonstrate 13C-methyl labeling of all these amino acids and additionally of alanine and phenylalanine. Alanine can be important as a more sensitive alternative to 15N-labeling, as it can also be considered a probe of the protein backbone. The named amino acids can all be labeled with our general protocol. First, amino acid specific labeling of Val and Leu will be shown, and then Ala, Met and Ile will be used to illustrate special cases, like suppression of unwanted metabolism, labeling in full media, and spectroscopic techniques to obtain high resolution spectra even if selectively methyl-labeled amino acids are not available.

Methyl group labeling of Val and Leu was achieved using 150 mg/l of 13Cγ-valine and 230 mg/l of 13Cδ-leucine (Tracer Technologies, Inc.), and 13C-incorporation of 60% and 71% was achieved, respectively. Alternatively, for these amino acids, an approach based on more economic precursors works in mammalian cells. For Val and Leu, the precursors α- keto-isovalerate and α-keto-isocaproate[44–46], respectively, are readily converted to the corresponding amino acids Val and Leu by HEK cells. Using precursors allows accessing local deuteration in a relatively inexpensive manner. As shown in Figure 4, specific deuteration leads to markedly sharper signals for Val and Leu methyl groups[47]. However, here, some scrambling occurs: α-keto-isovalerate is also incorporated into leucine residues, albeit at a much lower degree than into valine. This can be prevented by adding an excess of unlabeled leucine, which shifts the metabolic equilibria towards valine. Employing 160 mg/l of 2H,13C α-keto-isovalerate (L03, MagLab) and 400 mg/l of unlabeled leucine resulted in essentially scrambling-free spectra. For labelling of leucine, 160 mg/l of 2H,13C α-keto-isocaproate (L05, MagLab) were employed and no significant scrambling was observed. Also here, V3⊖ medium was used as the base of the medium and unlabeled amino acids missing from the omitted yeast extracts were compensated using an amino acid mix. In Figure 4, spectra of 13C-methyl labeled mEGFP are summarized.

#### Special case I: Amino acid type specific labeling in full medium with amino acid in excess or by prior enzymatic depletion

In principle, isotope labeling can be achieved using full medium by adding the desired amino acid in ten-fold excess over the amount of unlabeled amino acid in the medium. For some amino acid types, where the isotope labelled form is available at comparatively low cost, like 13Cε methionine, this approach can be economic, since no custom medium is required. Addition of 500 mg/l of 13Cε methionine to V3 medium prior to induction, yielded an isotope incorporation of 94%.

A variant of this approach, is using the enzyme methionine-γ-lyase (MGL) to degrade unlabeled methionine in the full medium – which in this case can be any medium formulation[48]. To this end, 2 mg/l MGL must be added to the medium, as well as the required co-factor pyridoxalphosphate (0.1 mM) and, depending on the media, additional buffering might be required. Thanks to volatile products of the reaction, this enzymatic reaction runs to completeness and after 48 h at 37°C the medium is devoid of any unlabeled methionine. Before addition of isotope labeled methionine, the MGL enzyme must be inhibited with 1 mM propargylglycine, which is well-tolerated by the cells. To demonstrate methionine labeling in other media with unknown composition, we performed a labeling experiment in HEK293 Freestyle medium (GIBCO Invitrogen), after treatment with MGL (Figure 5A).

**Figure 5:**
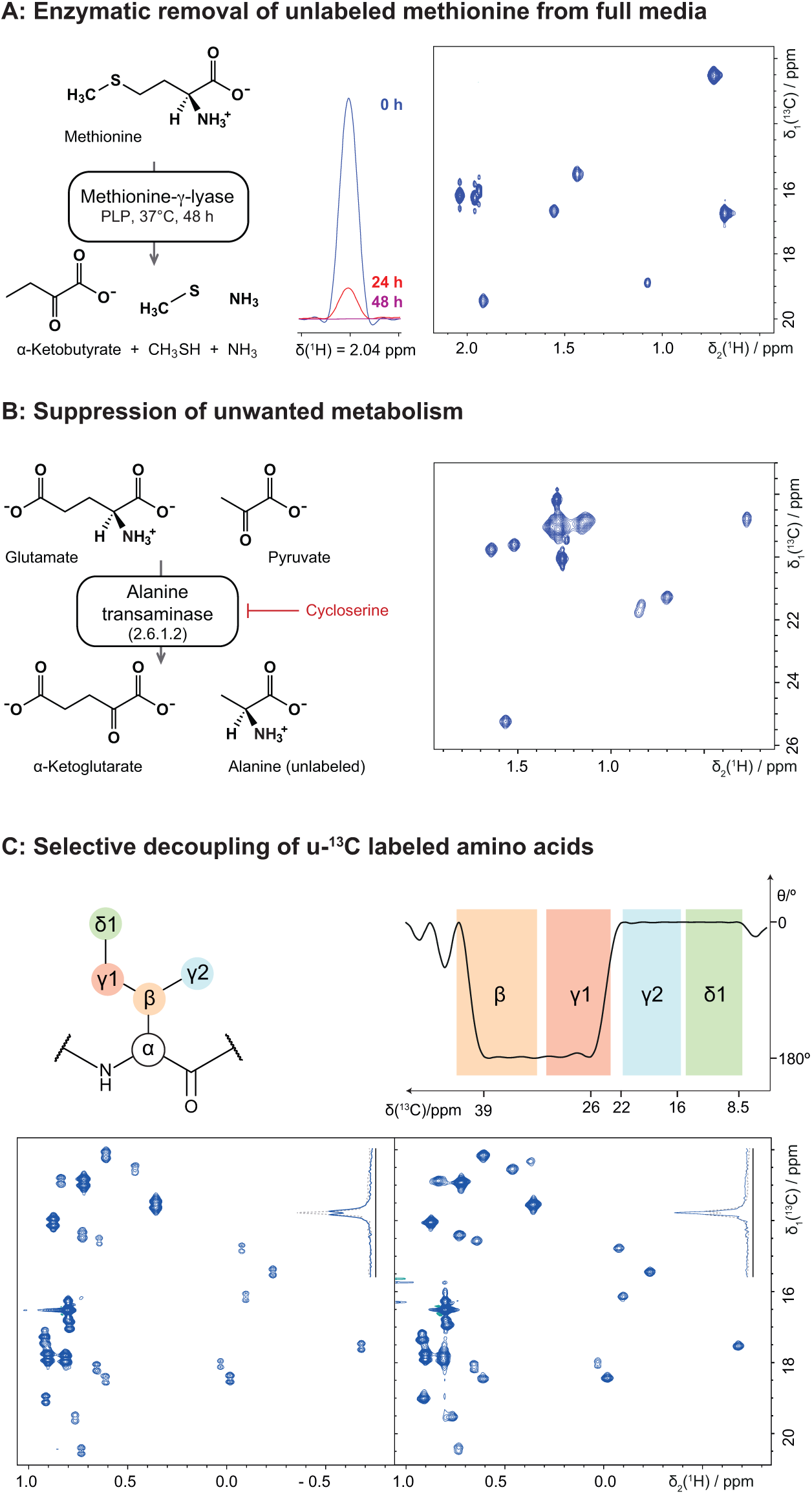
(A) Methionine labeling in full media by enzymatic removal of intrinsic methionine from a medium with unknown composition, here HEK293 Freestyle medium (GIBCO). On the left, the reaction of the enzyme is shown (for reaction conditions see main text). Note that methanethiol is volatile and thus is continuously removed from the reaction resulting in complete turnover of methionine. In the middle, the methionine signal in the medium is shown at 0, 24 and 48 h after treatment with MGL. To monitor the reaction, the medium was spiked with 1 g/l 13C-Met and 13C-edited 1D 1H spectra were recorded. On the right, a 2D [13C,1H] HMQC spectrum of 13C-Met labelled mEGFP is shown, produced from enzymatically methionine-free HEK293 Freestyle medium (GIBCO). (B) Isotope labeling of alanine with inhibition of alanine transaminase to suppress unwanted isotope dilution from pyruvate. On the left, the reaction of alanine transaminase is shown. Note that unlabeled pyruvate is present in rather high concentrations in the cell, thus driving the reaction towards the product alanine. Addition of L-cycloserine (red) inhibits this reaction. On the right, a 2D [13C,1H]-XL-ALSOFAST-HMQC spectrum of 2Hα,13Cβ-Ala labeled mEGFP is shown, which was produced in presence of 50 mg/l of L-cycloserine. (C) When selectively ^13^C methyl labeled amino acids are not available, suppression of ^1^*J*CC couplings in u-^13^C labelled amino acids yields sharp signals. On the top left, a schematic representation of isoleucine is shown and on the right, the Mz- excitation profile of an adapted BADCOP1 pulse is shown. The pulse was time-reversed and the length adjusted in order to selectively invert β and γ1 nuclei, resulting in decoupling of the adjacent γ2 and δ1 nuclei, respectively. (BADCOP1_TR, pulse length: 1700 μs, offset: 16 ppm, simulated in Topspin shapetool). Typical chemical shifts of carbon nuclei are indicated with colored bands (BMRB, [49]). Below 2D [13C,1H]-XL-ALSOFAST-HMQC spectra are plotted, showing the methyl region of 13C-Ile labeled mEGFP, without (left) and with (right) application of a BADCOP1_TR 180° pulse. A slice through one representative signal is shown in the top right corner of each spectrum and illustrates the suppression of the homonuclear couplings as well as the concurrent ∼2-fold gain in sensitivity. Spectra were recorded on Bruker Avance IIIHD 800 MHz (A) and 900 MHz (B, C) spectrometers equipped with TCI CryoProbes.

#### Special case II: Metabolism of human cells leading to isotope dilution

The amino acid metabolism of human cells has only low activity. One example shown above was the unwanted incorporation of α-keto-isovalerate into leucine, which could be controlled by shifting the metabolic equilibrium by adding an excess of unlabeled leucine. Another metabolic activity often observed in cell culture is the Cahill cycle [50], in which cells produce large amounts of unlabeled alanine from pyruvate, a highly abundant metabolite stemming from glycolysis (Figure 5B). In order to obtain sufficiently high incorporation of e.g. 13C-Ala, the enzyme alanine-transaminase, which is responsible for this reaction must be inhibited, which is easily achieved by adding 50 mg/l of L-cycloserine to the medium [14,51]. In our HEK293 cultures in V3 medium, we did not observe the detrimental levels of alanine production as for example previously in insect cells, but as a precaution, cycloserine was included in alanine labeling experiments, since it had no apparent negative effect on protein yields.

With this procedure a labeling efficiency of 87% could be achieved for 2Hα, 13Cβ-Ala, when employing 250 mg/l of labeled alanine (Figure 5B).

#### Special case III: Spectroscopic suppression of 1*J*_CC_ couplings for u-13C-labeled amino acids

For recording NMR spectra of methyl groups of proteins, amino acids with only the methyl carbon 13C-labelled are highly preferred over uniform 13C-labelled amino acids, because otherwise 1*J*CC couplings from the neighboring 13C nucleus lead to splitting of the signals and thereby limit the resolution to 120–150 Hz in the carbon dimension. To avoid split signals, either the approach using selectively 13C-methyl labelled *E. coli* extracts can be used or amino acids with this labeling pattern need to be purchased. Currently only selected amino acids like Ala, Met, Val and Leu are available with specific 13C-labeling of methyl groups from vendors and some only at high expense [47]. In contrast all amino acids can be commercially acquired in uniformly 13C-labeled form (u-13C).

Here, modified spectroscopic techniques set the path for obtaining spectra with high resolution from uniformly 13C labeled amino acids. The recently introduced BADCOP pulses based on optimal control, are highly band selective and therefore allow differential treatment of methyl group carbons and their adjacent carbon[52]. Time-reversal of the BADCOP1 pulse and adjustment of its duration allowed selective decoupling of the γ2 and δ1 methyl groups in Ile from β and γ1 carbons, respectively, yielding high resolution spectra (Figure 5C). In this specific case, spectra were recorded at 900 MHz using a BADCOP1_TR pulse of 1700 μs duration and 16 ppm offset. The protein was isotope labeled employing 250 mg/l of 13C6 Ile and 73% incorporation was achieved.

#### Choice of labeling pattern: spectral quality and coverage

The methods developed here enable studies of biologically and pharmaceutically highly relevant proteins that were previously not amenable to NMR. In Figure 7, a number of examples from our projects are compiled, namely the GPCRs bovine rhodopsin, human β1 and β2 adrenergic receptors and the transcription factor ERRγ. For all proteins, expression in HEK cells was rather straightforward, as determined in small scale expression test. However, we first wanted to determine the most suitable isotope pattern for these relatively large and typically flexible proteins. To this end, we used the stabilized β1 adrenergic receptor from turkey (*mg*β1AR, [53]) as a stepping stone to explore different isotope labeling patterns (13Cβ-Ala, 13Cγ-Val, 13Cδ-Leu, u-13C-Ile, 13Cε-Met). From the spectra shown in Figure 6, average signal intensities could be extracted, which established the following ranking for signal intensity: 13Cε-Met > 13Cδ1-Ile > 13Cδ-Leu ≈ 13Cγ-Val > 13Cβ-Ala. Additionally evaluating signal overlap, it is evident that 13Cε-Met yields spectra of highest quality. When choosing the labeling pattern, however, coverage of important sites of a protein needs to be considered as well. In this respect, methionine yields the poorest coverage, as it is the least abundant of all methyl-bearing amino acids. Depending on the amino acid composition of the target protein, or the site of interest, other amino acids might be more suitable. Another aspect regarding coverage, is whether side chain or backbone reporters are wanted. For the latter, 13Cβ-Ala is ideally suited, as the methyl groups is firmly attached to the backbone and has no rotational freedom with respect to backbone atoms. In fact, 13Cβ-Ala labelled receptor yielded high quality spectra as well, but overlap was already considerable. Modern algorithms capable of resolving overlap might therefore render Ala labeling an interesting alternative to amide labeling for monitoring backbone structure and dynamics.

**Figure 6:**
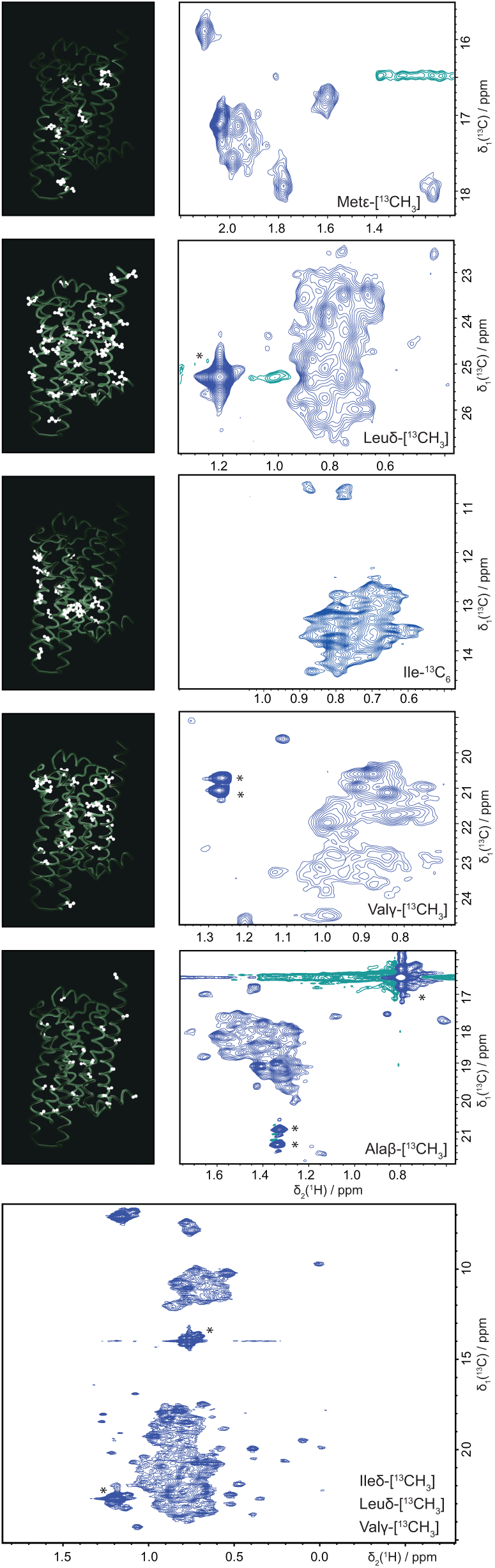
Choice of isotope labeling patterns based on coverage of the target protein (left) and quality of NMR spectra (right). On the left, ribbon representation of the GPCR β1AR of turkey (*mg*β1AR) is shown with amino acid side chains acting as small light sources illuminating parts of the protein. The illuminated parts should approximately represent the regions of the protein on which the labeled nuclei can report, as their resonance is influenced by effects in their immediate surroundings. On the right, [^13^C,^1^H]-XL-ALSOFAST-HMQC spectra of *mg*β1AR are shown, with ^13^C methyl group labeling of Met, Leu, Ile, Val, Ala and ILV as indicated in the spectrum. All spectra were recorded at 298 K on Bruker Avance IIIHD 900 MHz and NEO 1.2 GHz (bottom spectrum) spectrometers equipped with a TCI CryoProbes.

**Figure 7:**
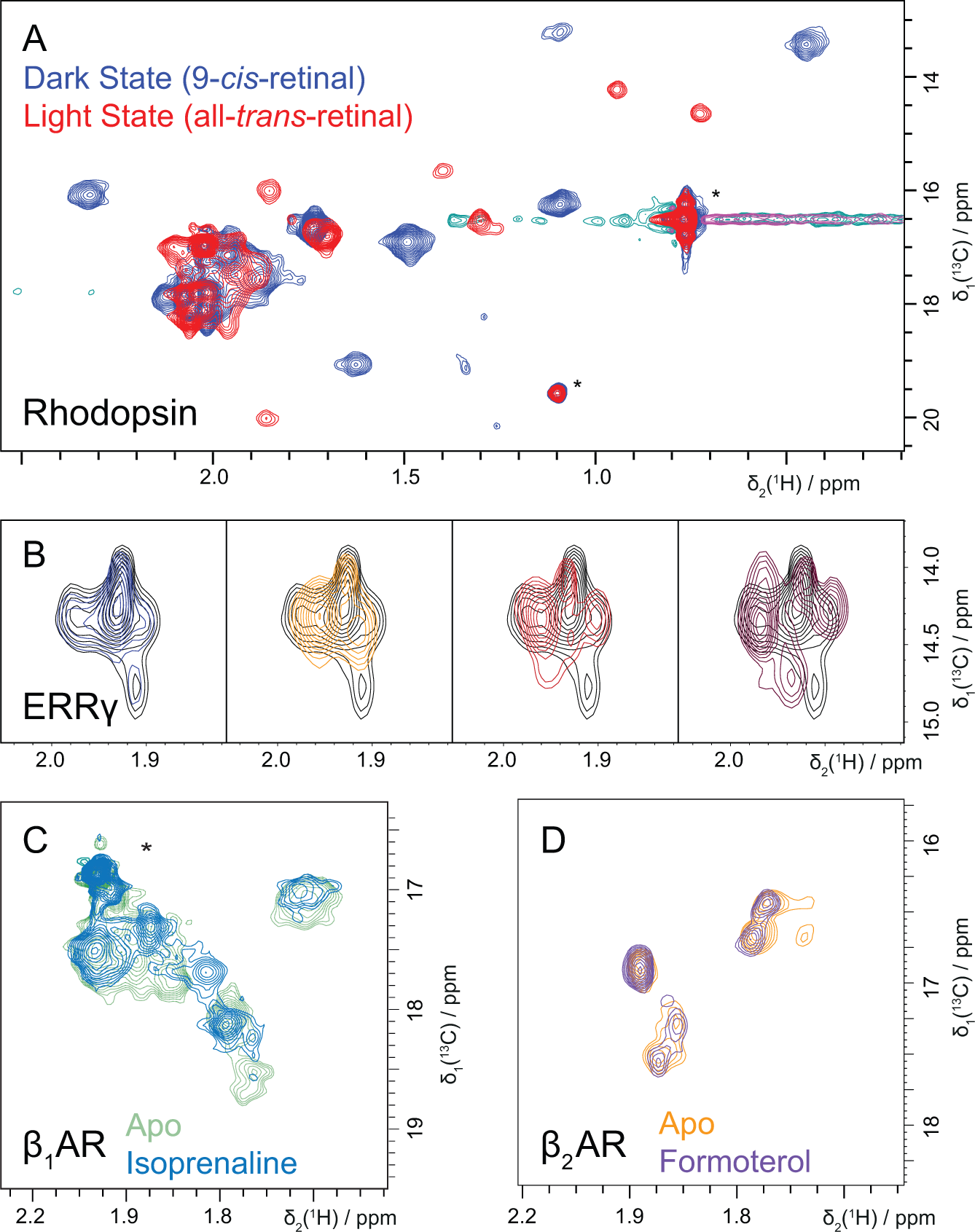
Examples of NMR studies enabled by mammalian isotope labeling. (A) Overlay of [13C,1H]-XL-ALSOFAST-HMQC spectra of bovine rhodopsin in DM micelles (0.2%) in its dark (blue) and light-activated state (red). NMR spectra of the stable dark state protein were recorded in 3 mm NMR tubes, which were protected from light by coverage in electric heat shrink. Light state spectra were obtained after 1 min illumination of the sample with a flash light of a mobile phone. The large chemical shift changes affecting essentially all resonances reflect the global conformational change of the receptor in response to the isomerization of 9-*cis* to all-*trans* retinal. (B) Overlay of [13C,1H]-HMQC spectra of estrogen related receptor γ (ERRγ) in absence (black contour lines) and presence of different ligands (contour lines of different colours). These simple spectra show binding of all compounds except for the one giving rise to the blue spectrum on the left-most spectrum. (C) Overlay of [^13^C,^1^H]-XL-ALSOFAST-HMQC spectra of stabilized human β1 adrenergic receptor (*hs*β1AR) in LMNG/CHS (0.01/0.001%) micelles without (blue) and with (green) the agonist isoprenaline bound. Also here, chemical shift changes reflect the change in equilibrium towards an active conformation upon agonist binding. (D) [13C,1H]-XL-ALSOFAST-HMQC spectrum for human β2 adrenergic receptor (*hs*β2AR) in MSPΔH5 nanodiscs (PO/PG) in its resting state (orange) and in presence of 1 mM Formoterol (violet). Spectra were recorded on Bruker Avance IIIHD 600 MHz (B) and 900 MHz (A, C, D) spectrometers equipped with TCI CryoProbes.

For the receptors at hand, 13C-methyl labeling of methionine seemed to be the most viable option, since methionine residues are quite evenly spread over the protein, and 13C- labeled methyl groups of Val, Leu and Ile yielded spectra with strong overlap of exchange broadened signals, emphasizing the challenges that these highly flexible family of receptors pose to NMR studies.

#### Enabling aspects of isotope labeling in mammalian cells

For all the following examples we used 13Cε-Met labeling, because of the superior spectral quality. First, we present bovine rhodopsin, for which expression in HEK293 cells was previously established for the constitutively active variant (N2C, M257Y, D282C) [54–56], and 13Cε-Met isotope labeling was thus straightforward. Commercially available 9- *cis*-retinal was used as surrogate for the natural ligand 11-*cis*-retinal. The obtained protein preparation in DM micelles was tested by UV-visible spectroscopy to test for light contamination and to assess the quality of the sample. Functional integrity of the produced protein was demonstrated by recording spectra of the dark state of rhodopsin and of the same protein after illumination. Essentially all of the visible resonances of the receptor shift upon illumination, reflecting the overall conformational change of rhodopsin, when changing from an inactive state to an activated state. Rhodopsin is exemplary for a multitude of proteins, which require a higher expression host, and where production in mammalian cells is already established. Thanks to the above presented methods such proteins are now readily amenable to NMR studies.

Pharmaceutically highly relevant targets like GPCRs can be studied by NMR [57–60]. However, establishing a single receptor for NMR characterization is already a magnificent task and therefore even large NMR laboratories typically work on just one specific type of receptor. Since HEK293 cells are highly competent at expressing receptors, we were able to not only produce the above-mentioned rhodopsin in labeled form, but also the pharmacologically relevant human β1 and β2 adrenergic receptors, which are the target of some of the most prescribed drugs.

For producing human β1 and β2 adrenergic receptors (*hs*β1AR and *hs*β2AR, respectively) we therefore focused on 13Cε-Met labeling. For *hs*β1AR we transferred the mutations found to stabilize *mg*β1AR [60] to the human counterpart and were able to obtain NMR spectra of higher quality than with wt protein. The construct can be activated by agonists and binds to G-protein surrogates. NMR studies of human β1AR have not been reported before, and to our knowledge the spectra in Figure 7 are the first ones published. Our laboratory was able to produce these different receptors with relative ease. We attribute this to expression in mammalian cells. Since currently HEK293 cells are probably the most competent expression host for human proteins, the first step in protein production – expression of correctly folded protein – is solved, and one can focus on optimization of subsequent steps.

A further example of application of labeling in mammalian cells enabling medicinal research is the estrogen-related receptor γ (ERRγ) [61]: This difficult-to-produce protein was labeled in mammalian cells and could be used to test ligands discovered in biochemical assays for binding. In Figure 7 a number of successful ligand binding experiments are shown, where the ligands induced chemical shift perturbations, which are indicative of binding.

In summary, we show that with the ability of performing isotope labeling in mammalian cells, a large hurdle towards NMR studies of difficult-to-produce proteins is removed. Sufficient amounts of protein can be produced and isotope incorporation is high enough for functional studies and for characterizing molecular interactions at the atomic level.

## Methods

For better readability, the protocols for isotope labeling can be found in a separate file in the supplementary information. Here, purification protocols for individual proteins are given.

Purification of *hs*β1AR, *hs*β2AR and *mg*β1AR: The frozen cell pellet was resuspended in solubilisation buffer (20 mM HEPES (pH 7.5), 300 mM NaCl, 1 mM EDTA, 10% glycerol, 1 mM PMSF and 1% LMNG (w/v, *hs*β1AR), 2% DM (w/v, *mg*β1AR) or 1% DDM (w/v, *hs*β2AR)) and treated for 20 s with a TURRAX IKA T18 disperser equipped with a S18N- 19G dispersing tool. The homogenised solution was stirred at 4°C for 1 h and subsequently cleared by centrifugation at 185,000 g for 1 h. The supernatant was filtered using a 0.45 μm nitrocellulose membrane (Merck, MF-Millipore) and subsequently loaded onto a 5 ml StrepTrap HP column (GE). The column was washed with 10 column volumes of washing buffer (20 mM HEPES pH 7.5, 300 mM NaCl, 0.01% LMNG (w/v) and 0.001% CHS (w/v), or 0.1 % DDM (w/v) or 0.2% DM (w/v)) and afterwards protein was eluted with 4 column volumes of elution buffer (20 mM HEPES pH 7.5, 300 mM NaCl, 2.5 mM desthiobiotin, 0.01% LMNG (w/v) and 0.001% CHS (w/v), or 0.1 % DDM (w/v) or 0.2% DM (w/v)). 3C HRV protease and 1 mM DTT were added to the eluted protein and incubated overnight at 4°C to cleave off the mEGFP-tag.

Afterwards, the protein was concentrated in a 50 kDa concentrator and further purified on a Superdex 200 column (GE) for size-exclusion chromatography in SEC buffer (20 mM HEPES pH 7.5, 300 mM NaCl, 0.01% LMNG (w/v) and 0.001% CHS (w/v), or 0.1 % DDM (w/v) or 0.2% DM (w/v)). For *hs*β1AR and *mg*β1AR, the receptor-containing fractions were pooled and concentrated in a 50 kDa concentrator (Amicon Ultra, Merck MF-Millipore) to a concentration above 50 μM and subsequently diluted with D2O and SEC buffer to 50 μM with 10% D2O. These samples were aliquoted in 100μl fractions and flash frozen in liquid nitrogen. Aliquots were stored at −80°C prior to use. For *hs*β2AR the SEC eluate was only concentrated to 700 μL and further transferred into nanodiscs. To this end 100 mM stock solutions of POPC and POPG powders in 200 mM sodium cholate were prepared by repeated shock-freezing of the suspension in liquid nitrogen and subsequent thawing at RT, finally resulting in a clear solution. To 700 μL of *hs*β2AR soluiton, 60 μL POPC and 40 μL POPG stock solution were added and the sample was incubated for 5 min on ice. Subsequently, the sample was supplemented with MSP1D1ΔH5 and the volume was increase to 1 mL with SEC buffer to achieve final concentrations of 10 mM lipids (with a 3:2 ratio of POPC:POPG), 20 mM sodium cholate and 200 μM MSP1D1ΔH5. This mixture was incubated over night at 4 °C. 800 mg Bio-Beads (SM-2 resin, Bio-Rad) were added in 4 portions to the solution. After every addition the mixture was stored for 1.5 h on ice and mixed from time to time by inversion of the reaction tube. Bio-Beads were separated from the solution in Mobicols, using filters with 35 μm pore size (MoBiTec), by centrifugation for 1 min at 1500 g. Removal of tags was achieved by the addition of 50 μL 3C HRV protease (in-house production, His-tagged, 1 mg) as well as 1 mM DTT and incubation over night. Subsequently, the mixture was purified by SEC on a Superdex 200 Increase 10/300 column (GE) with SEC buffer (20 mM HEPES pH 7.5, 150 mM NaCl). Fractions with GPCR-containing nanodiscs were pooled and concentrated in 50 kDa Amicon-Ultra 0.5 mL centrifugal filters (Merck Millipore) to reach a final concentration of ∼60 μM. The sample was stored at 4 °C until further usage.

Bovine rhodopsin and MSP1D1ΔH5 were purified according to established protocols [56,62].

## Discussion

We present here a suite of protocols for isotope labeling in mammalian cells, which enables studying difficult proteins by NMR. Wherever possible, priority was given to reduce cost and lower the complexity of every step in the process, starting from simple laboratory equipment, over efficient suspension culture to a diverse set of labeling schemes which are highly cost-effective compared to current standards. This simple approach should lower the hurdle for biology-focused laboratories to produce isotope labeled proteins or, vice versa, for NMR laboratories to enter mammalian cell culture.

### General cell culturing setup for cost-effective isotope labeling

The choice of suspension culture in flasks over adherent culture in plates significantly reduces the workload and fits the relatively high protein amounts required for NMR studies better than currently established adherent culture.

Our protocols are compatible with both, transient and stable transfection of cell lines, thus offering flexibility to choose between fast turnover when expressing for instance different point mutants for assignment purposes, and reproducibly high yields for the core protein of a project, respectively.

For isotope labeling one needs to balance costs against label incorporation. Since the hurdle to enter mammalian cell culture is rather high, we emphasized economy over label incorporation, such that our protocols typically yield isotope incorporation levels in the order of 60–95%. Highest incorporation could be achieved by using media completely devoid of unlabeled amino acids (for which we found no vendor) and by culturing cells for several generations in labeled medium. In order to save costs, in our protocols we grow cells in unlabeled medium and only employ labeled medium for the actual expression. Thus, there is inevitable carry-over of unlabeled metabolites, lowering the final isotope incorporation by estimated 10–15%. Cells could be grown in labeled medium for 2–3 passages before expression in labeling medium, but the cost would consequently double or triple. Our approach thus essentially excludes efficient triple resonance experiments, but since most proteins requiring mammalian cells for their production are larger than 30 kDa, anyway other methods are employed for resonance assignment. 2D heteronuclear correlation experiments in turn still remain highly sensitive even with these slightly lower incorporation levels, and a the principal experiments used in functional NMR studies.

### Uniform labeling

For uniform labeling, mammalian cells require media containing all essential amino acids in isotope labeled form, and growth rates increase significantly if all amino acids are present in the medium. Since adding twenty individual isotope labeled amino acids to a medium is prohibitively expensive, we rely on labeled amino acid extracts from organisms that can grow in inexpensive minimal media, that is, yeast, single-celled algae or *E. coli* bacteria. With this approach uniform labeling becomes ∼25-fold less expensive compared to using individual amino acids, and 10-fold less expensive than using the available commercial medium for uniform 15N-labeling in mammalian cells. On top of that one should mind that the commercial medium can only be used in adherent culture, such that one needs to factor in the additional labor for expanding cells to hundred plates, which takes weeks, and then transfecting, inducing, and finally scratching such a number of plates. Therefore, in summary, using labeled amino acid mixtures in suspension culture massively simplifies expression of uniformly labeled proteins and reduces costs at the same time.

Amino acid extracts can have different characteristics depending on the type of hydrolysis the extracts were subjected [63]. We mainly distinguish acid hydrolysis and autolysis, where the former is a rather harsh method where Asn and Gln side chain amides are hydrolyzed to the respective acids, Asp and Glu, and Trp and Cys are typically oxidized to different products. Enzymatic autolysis by intrinsic proteases results in autolysates which typically conserve all amino acids. The presence of Asn/Gln and Trp can easily be checked by 2D [15N,1H]-HSQC experiments on 15N-labeled extracts, where the distinct side chain amide signals can be identified. Asn and Gln are not essential amino acids for mammalian cells, but cell culture parameters are typically better if especially Gln in present in the medium. Therefore, whenever possible, we prefer extracts based on enzymatic lysis, which conserves all amino acids.

We showed that crude extracts from different organisms (yeast, algae, bacteria) can be used for the purpose of isotope labeling. The different extracts have their respective advantages and disadvantages. Based on our results that about 40% higher yields are obtained using yeast extracts over algal extracts, we suggest that for uniform 15N labeling, yeast extracts should be employed and for other uniform labeling patterns, algal extracts should be used, because for 13C and 2H labeling, they are much more economic, even when doubling the culture volume is required (to our knowledge, 13C- and 2H-labeled yeast extracts are not available and if they were, they would be extremely expensive due to the high cost of labeled glucose required as starting material). Extracts from *E. coli* make more complex labeling patterns accessible, as demonstrated for ILV 13C -methyl labeling, but typically will be the most expensive to produce. However, they might be a suitable source of labelled amino acids as a byproduct in some NMR laboratories, which produce ILV 13C-methyl labelled samples in *E. coli* on a regular basis and might otherwise discard cell pellets after purification of their target protein. Thus, in summary, we suggest yeast extracts for u-15N labeling, algal extracts for u-13C and *E. coli* for special labeling patterns that can’t easily be produced in any of the other organisms.

In this work, we emphasized uniform 15N and 13C labeling and deuteration was not a major focus. Since reaching the high incorporation levels required for triple resonance experiments is hardly feasible, there is little incentive to attempt uniform deuteration. Where we see the highest potential for 2H-labeling is in specific deuteration of selected amino acids as exemplified with Ala, Val and Ile. Arthanari and co-workers have demonstrated fantastic signal improvement by selective deuteration of individual sites in close proximity of observed groups [47], and this can be achieved best with our protocols for amino acid specific labeling, provided the labeling pattern is available for a given amino acid. For such a labeled amino acid the local incorporation is therefore 100%, which leads to an ideal situation for spectroscopy. Thus, in the following we turn to amino acid type specific labeling.

### Amino acid type specific labeling

For amino acid type specific labeling, there are only few limitations. Since amino acid metabolism in HEK293 cells is much lower than e.g. in *E. coli*, isotope labels are rarely transferred to other amino acids (label scrambling) and endogenous production of unlabeled amino acids (label dilution) is limited to few well-understood cases. Scrambling of the label was hardly ever observed, due to low amino acid metabolism of HEK293 cells. Only for 15N-Val, minimal scrambling of the 15N isotope was observed. For all 13C-labeling patterns we tested, scrambling could only be detected for α-keto- isovalerate to leucine. Based on our experience with insect cells, otherwise we expect only the amino acids connected to the central citric acid cycle to be difficult to label, that is Asn and Gln, and to a lower extent also Asp and Glu. Isotope dilution was only detected for 13C-Ala labeling, where isotope dilution from unlabeled pyruvate was observed, but this unwanted reaction could be inhibited by the use of the specific inhibitor cycloserine. One limitation is that HEK293 cells typically require the entire amino acid or a very close precursor in the growth medium. Therefore, full amino acids with the desired labeling pattern are required. Currently, it is possible to obtain every amino acid in uniformly labeled form, however, specific labels, like for example the highly sought 13C-methyl variants of amino acids are rare, especially including deuteration. We presented a spectroscopic solution for working with uniformly 13C-labeled amino acids and were able to obtain decoupled spectra of methyl groups of u-13C-Ile with high intensity. This alleviates the problem, but at the same time we trust that further developments in amino acid synthesis will make specifically 13C-methyl labeled amino acids available in the near future, since demand is clearly increasing.

Currently, the demand for specifically labeled amino acids is rising due to the increasing use of insect cell culture in the community. Isotope labeling in insect cells has been established in several laboratories and is the main eukaryotic system used for heterologous expression for NMR studies. Therefore, a comparison of insect and mammalian expression systems for producing isotope labeled proteins is important. Laboratory requirements for the two are very similar, except that human cells require an incubator with controlled CO2 levels and humidity. Media for both types of cells are similar in the ingredients, however, insect cell media contain 10-fold higher amounts of amino acids. Analysis of insect cell media reveal a total of 15 g of amino acids per liter (e.g. ExCell 420), while the HEK293 media used here only contain about 1.5 g of total amino acids, with a significant potential for lower costs. In practice however, for uniform isotope labeling, in insect cells only 10 g/l of algal extracts are employed, in human cells 4 g of algal or 5 g of yeast extracts, thus reducing the isotope consumption only by a factor of 2. Therefore, it probably depends on the expression yields of every given protein construct, whether insect or mammalian cells are more economic, however, with mammalian cells being generally less isotope-consuming. In terms of workload, transient expression in human cells is most efficient, since preparation of DNA requires just 2–3 days. For insect cells, a sufficient quantity of baculovirus stock first needs to be generated, which typically takes one month. Finally, selection of a stable human cell line and adaptation to suspension culture requires 1–2 months. Thus, mammalian cells are the more efficient system for screening several constructs, for preparation of a long-lasting expression system both organisms are comparable. Again it will depend on the individual target protein, whether the advantage is with insect or mammalian cell culture. Here, if an overall trend should be identified, it appears that cytosolic proteins have higher expression levels in insect cells, while membrane proteins and secreted proteins typically have higher yields in mammalian cells. The most impressive example are industrially produced therapeutic antibodies, which are often expressed at more than 1 g/l of cell culture.

In summary, the methods presented here, enable a wide range of labeling patterns with stable isotopes in HEK293 cells, which represent the most fitting expression system for human proteins. Here, the chances of successful expression of difficult proteins are highest compared to bacterial and insect cells, making all classes of human proteins more accessible. The hurdles to establish a eukaryotic expression system in an NMR laboratory were high, regarding both cost and work. Preparing a 15N-labeled sample – the only commercially available labeling pattern – signified expanding an adherent culture to dozens of plates, impractical transfection of all those plates and finally scratching them at harvest and at the same time spending thousands of EUR on medium. We therefore shed costs in every step and simplified protocols considerably, resulting in a simple setup (shaking plate for suspension culture in regular incubator) and an order of magnitude lowered costs for media, and at the same time higher resulting protein amounts and much wider range of accessible labeling patterns (u-15N, u-13C, amino acid-type selective labeling, methyl-group labeling).

We therefore hope that these methods signify a breakthrough allowing to fully exploit the high information content of NMR for biological and biomedical research.

## Acknowledgements

The authors would like to acknowledge Chia-wei Tan-Lin and Paolo Nanni from the functional genomics center Zürich for excellent help with mass spectrometry. P.R., A.L. and L.R.F. thank the Biomolecular Structure and Mechanism PhD Program of the Life Science Zurich Graduate School. This work was supported by the Swiss National Science Foundation by grants 31-179319 and 31-208029 to A.D.G.

## Supplementary material

Detailed protocols are available as supplementary material.

## References

[1] T.L. Blundell, H. Jhoti, C. Abell, High-Throughput Crystallography for Lead Discovery in Drug Design, Nature Reviews Drug Discovery. 1 (2002) 45–54. 10.1038/nrd706.

[2] M. Pellecchia, I. Bertini, D. Cowburn, C. Dalvit, E. Giralt, W. Jahnke, T.L. James, S.W. Homans, H. Kessler, C. Luchinat, Perspectives on NMR in drug discovery: a technique comes of age, Nature Reviews Drug Discovery. 7 (2008) 738–745.

[3] W. Kühlbrandt, The Resolution Revolution, Science. 343 (2014) 1443. DOI: 10.1126/science.125165.

[4] X. Bai, G. McMullan, S.H.W. Scheres, How cryo-EM is revolutionizing structural biology, Trends in Biochemical Sciences. 40 (2015) 49–57. 10.1016/j.tibs.2014.10.005.

[5] C.A. Lutomski, T.J. El-Baba, C.V. Robinson, R. Riek, S.H.W. Scheres, N. Yan, M. AlQuraishi, L. Gan, The next decade of protein structure, Cell. 185 (2022) 2617–2620. 10.1016/j.cell.2022.06.011.

[6] I. Jelesarov, H.R. Bosshard, Isothermal titration calorimetry and differential scanning calorimetry as complementary tools to investigate the energetics of biomolecular recognition, J. Mol. Recognit. 12 (1999) 3–18. 10.1002/(SICI)1099-1352(199901/02)12:1<3::AID-JMR441>3.0.CO;2-6.

[7] C. Boozer, G. Kim, S. Cong, H. Guan, T. Londergan, Looking towards label-free biomolecular interaction analysis in a high-throughput format: a review of new surface plasmon resonance technologies, Current Opinion in Biotechnology. 17 (2006) 400–405. 10.1016/j.copbio.2006.06.012.

[8] R. Assenberg, P.T. Wan, S. Geisse, L.M. Mayr, Advances in recombinant protein expression for use in pharmaceutical research, Current Opinion in Structural Biology. 23 (2013) 393–402. 10.1016/j.sbi.2013.03.008.

[9] F.J. Fernández, M.C. Vega, Technologies to keep an eye on: alternative hosts for protein production in structural biology, Current Opinion in Structural Biology. 23 (2013) 365–373. 10.1016/j.sbi.2013.02.002.

[10] S. Gräslund, P. Nordlund, J. Weigelt, J. Bray, O. Gileadi, S. Knapp, U. Oppermann, C. Arrowsmith, R. Hui, J. Ming, S. dhe-Paganon, H. Park, A. Savchenko, A. Yee, A. Edwards, R. Vincentelli, C. Cambillau, R. Kim, S.-H. Kim, Z. Rao, Y. Shi, T.C. Terwilliger, C.-Y. Kim, L.-W. Hung, G.S. Waldo, Y. Peleg, S. Albeck, T. Unger, O. Dym, J. Prilusky, J.L. Sussman, R.C. Stevens, S.A. Lesley, I.A. Wilson, A. Joachimiak, F. Collart, I. Dementieva, M.I. Donnelly, W.H. Eschenfeldt, Y. Kim, L. Stols, R. Wu, M. Zhou, S.K. Burley, J.S. Emtage, J.M. Sauder, D. Thompson, K. Bain, J. Luz, T. Gheyi, F. Zhang, S. Atwell, S.C. Almo, J.B. Bonanno, A. Fiser, S. Swaminathan, F.W. Studier, M.R. Chance, A. Sali, T.B. Acton, R. Xiao, L. Zhao, L.C. Ma, J.F. Hunt, L. Tong, K. Cunningham, M. Inouye, S. Anderson, H. Janjua, R. Shastry, C.K. Ho, D. Wang, H. Wang, M. Jiang, G.T. Montelione, D.I. Stuart, R.J. Owens, S. Daenke, A. Schütz, U. Heinemann, S. Yokoyama, K. Büssow, K.C. Gunsalus, Protein production and purification, Nature Methods. 5 (2008) 135–146. 10.1038/nmeth.f.202.

[11] A.M. Gronenborn, T. Polenova, Introduction: Biomolecular NMR Spectroscopy, Chem. Rev. 122 (2022) 9265–9266. 10.1021/acs.chemrev.2c00142.

[12] K. Wüthrich, NMR Studies of Structure and Function of Biological Macromolecules (Nobel Lecture), Angewandte Chemie International Edition. 42 (2003) 3340–3363. 10.1002/anie.200300595.

[13] A.D. Gossert, W. Jahnke, Isotope labeling in insect cells, in: H.S. Atreya (Ed.), Isotope Labeling in Biomolecular NMR, Springer, Dordrecht, 2012: pp. 179–196. http://www.springerlink.com/index/10.1007/978-94-007-4954-2 (accessed July 23, 2014).

[14] L. Skora, B. Shrestha, A.D. Gossert, Isotope Labeling of Proteins in Insect Cells, in: Z. Kelman (Ed.), Methods in Enzymology, Academic Press, Burlington, 2015: pp. 245–288. http://linkinghub.elsevier.com/retrieve/pii/S0076687915003110 (accessed November 23, 2015).

[15] B. Franke, C. Opitz, S. Isogai, A. Grahl, L. Delgado, A.D. Gossert, S. Grzesiek, Production of isotope-labeled proteins in insect cells for NMR., J Biomol NMR. 71 (2018) 173–184. 10.1007/s10858-018-0172-7.

[16] A. Strauss, F. Bitsch, B. Cutting, G. Fendrich, P. Graff, J. Liebetanz, M. Zurini, W. Jahnke, Amino–acid-type selective isotope labeling of proteins expressed in Baculovirus-infected insect cells useful for NMR studies, Journal of Biomolecular NMR. 26 (2003) 367–372. 10.1023/A:1024013111478.

[17] A. Strauss, F. Bitsch, G. Fendrich, P. Graff, R. Knecht, B. Meyhack, W. Jahnke, Efficient uniform isotope labeling of Abl kinase expressed in Baculovirus-infected insect cells, Journal of Biomolecular NMR. 31 (2005) 343–349. 10.1007/s10858-005-2451-3.

[18] K. Saxena, A. Dutta, J. Klein-Seetharaman, H. Schwalbe, Isotope Labeling in Insect Cells, in: A. Shekhtman, D.S. Burz (Eds.), Protein NMR Techniques, Humana Press, Totowa, NJ, 2012: pp. 37–54. http://www.springerlink.com/index/10.1007/978-1-61779-480-3_3 (accessed November 14, 2013).

[19] B. Jin, N. Thakur, A.V. Wijesekara, M.T. Eddy, Illuminating GPCR signaling mechanisms by NMR spectroscopy with stable-isotope labeled receptors, Current Opinion in Pharmacology. 72 (2023) 102364. 10.1016/j.coph.2023.102364.

[20] K. Büssow, Stable mammalian producer cell lines for structural biology, Current Opinion in Structural Biology. 32 (2015) 81–90. 10.1016/j.sbi.2015.03.002.

[21] F.L. Graham, Growth of 293 Cells in Suspension Culture, Journal of General Virology. 68 (1987) 937–940. 10.1099/0022-1317-68-3-937.

[22] M. Malm, R. Saghaleyni, M. Lundqvist, M. Giudici, V. Chotteau, R. Field, P.G. Varley, D. Hatton, L. Grassi, T. Svensson, J. Nielsen, J. Rockberg, Evolution from adherent to suspension: systems biology of HEK293 cell line development, Sci Rep. 10 (2020) 18996. 10.1038/s41598-020-76137-8.

[23] A. Dutta, K. Saxena, H. Schwalbe, J. Klein-Seetharaman, Isotope Labeling in Mammalian Cells, in: A. Shekhtman, D.S. Burz (Eds.), Protein NMR Techniques, Humana Press, Totowa, NJ, 2012: pp. 55–69. http://www.springerlink.com/index/10.1007/978-1-61779-480-3_4 (accessed November 14, 2013).

[24] A.P. Hansen, A.M. Petros, A.P. Mazar, T.M. Pederson, A. Rueter, S.W. Fesik, A practical method for uniform isotopic labeling of recombinant proteins in mammalian cells, Biochemistry. 31 (1992) 12713–12718. 10.1021/bi00166a001.

[25] D. Skelton, A. Goodyear, D. Ni, W.J. Walton, M. Rolle, J.T. Hare, T.M. Logan, Enhanced production and isotope enrichment of recombinant glycoproteins produced in cultured mammalian cells, Journal of Biomolecular NMR. 48 (2010) 93–102. 10.1007/s10858-010-9440-x.

[26] E. Luchinat, L. Banci, In-cell NMR: a topical review, IUCrJ. 4 (2017) 108–118. 10.1107/S2052252516020625.

[27] K. Werner, C. Richter, J. Klein-Seetharaman, H. Schwalbe, Isotope labeling of mammalian GPCRs in HEK293 cells and characterization of the C-terminus of bovine rhodopsin by high resolution liquid NMR spectroscopy, Journal of Biomolecular NMR. 40 (2008) 49–53. 10.1007/s10858-007-9205-3.

[28] T.A. Egorova-Zachernyuk, G.J.C.G.M. Bosman, W.J. DeGrip, Uniform stable-isotope labeling in mammalian cells: formulation of a cost-effective culture medium, Applied Microbiology and Biotechnology. 89 (2011) 397–406. 10.1007/s00253-010-2896-5.

[29] L. Barbieri, E. Luchinat, L. Banci, Characterization of proteins by in-cell NMR spectroscopy in cultured mammalian cells, Nat Protoc. 11 (2016) 1101–1111. 10.1038/nprot.2016.061.

[30] S. Yanaka, H. Yagi, R. Yogo, M. Onitsuka, K. Kato, Glutamine-free mammalian expression of recombinant glycoproteins with uniform isotope labeling: an application for NMR analysis of pharmaceutically relevant Fc glycoforms of human immunoglobulin G1, J Biomol NMR. 76 (2022) 17–22. 10.1007/s10858-021-00387-5.

[31] R.V. Williams, M.J. Rogals, A. Eletsky, C. Huang, L.C. Morris, K.W. Moremen, J.H. Prestegard, AssignSLP_GUI, a software tool exploiting AI for NMR resonance assignment of sparsely labeled proteins, Journal of Magnetic Resonance. 345 (2022) 107336. 10.1016/j.jmr.2022.107336.

[32] V. Tugarinov, L.E. Kay, 1H, 13C-1H, 1H dipolar cross-correlated spin relaxation in methyl groups, Journal of Biomolecular NMR. 29 (2004) 369–376.

[33] S. Schütz, R. Sprangers, Methyl TROSY spectroscopy: A versatile NMR approach to study challenging biological systems, Progress in Nuclear Magnetic Resonance Spectroscopy. 116 (2020) 56–84. 10.1016/j.pnmrs.2019.09.004.

[34] P. Rößler, D. Mathieu, A.D. Gossert, Enabling NMR Studies of High Molecular Weight Systems Without the Need for Deuteration: The XL-ALSOFAST Experiment with Delayed Decoupling, Angew. Chem. Int. Ed. 59 (2020) 19329–19337. 10.1002/anie.202007715.

[35] T. Yao, Y. Asayama, Animal-cell culture media: History, characteristics, and current issues, Reprod Med Biol. 16 (2017) 99–117. 10.1002/rmb2.12024.

[36] R.G. Ham, Clonal growth of mammalian cells in a chemically defined, synthetic medium, Proc Natl Acad Sci U S A. 53 (1965) 288–293. 10.1073/pnas.53.2.288.

[37] R. Dulbecco, G. Freeman, Plaque production by the polyoma virus, Virology. 8 (1959) 396–397. 10.1016/0042-6822(59)90043-1.

[38] B.P. Cormack, R.H. Valdivia, S. Falkow, FACS-optimized mutants of the green fluorescent protein (GFP), Gene. 173 (1996) 33–38. 10.1016/0378-1119(95)00685-0.

[39] D.A. Zacharias, J.D. Violin, A.C. Newton, R.Y. Tsien, Partitioning of Lipid-Modified Monomeric GFPs into Membrane Microdomains of Live Cells, Science. 296 (2002) 913– 916. 10.1126/science.1068539.

[40] E.G. Bligh, W.J. Dyer, A Rapid Method of Total Lipid Extraction and Purification, Can. J. Biochem. Physiol. 37 (1959) 911–917. doi.org/10.1139/o59-099.

[41] D.C. White, Evaluation of a hexane/isopropanol lipid solvent system for analysis of bacterial phospholipids and application to chloroform-soluble Nuclepore (polycarbonate) membranes with retained bacteria, J. Microbial Meth. 8 (1988) 131– 137.

[42] D.C. White, W.M. Davis, J.S. Nickels, J.D. King, R.J. Bobbie, Determination of the sedimentary microbial biomass by extractible lipid phosphate, Oecologia. 40 (1979) 51–62. 10.1007/BF00388810.

[43] R. Linser, V. Gelev, F. Hagn, H. Arthanari, S.G. Hyberts, G. Wagner, Selective Methyl Labeling of Eukaryotic Membrane Proteins Using Cell-Free Expression, Journal of the American Chemical Society. 136 (2014) 11308–11310. 10.1021/ja504791j.

[44] M.K. Rosen, K.H. Gardner, R.C. Willis, W.E. Parris, T. Pawson, L.E. Kay, Selective methyl group protonation of perdeuterated proteins, Journal of Molecular Biology. 263 (1996) 627–636.

[45] R.J. Lichtenecker, N. Coudevylle, R. Konrat, W. Schmid, Selective Isotope Labelling of Leucine Residues by Using α-Ketoacid Precursor Compounds, ChemBioChem. 14 (2013) 818–821. 10.1002/cbic.201200737.

[46] R. Lichtenecker, M.L. Ludwiczek, W. Schmid, R. Konrat, Simplification of Protein NOESY Spectra Using Bioorganic Precursor Synthesis and NMR Spectral Editing, J. Am. Chem. Soc. 126 (2004) 5348–5349. 10.1021/ja049679n.

[47] A. Dubey, N. Stoyanov, T. Viennet, S. Chhabra, S. Elter, J. Borggräfe, A. Viegas, R.P. Nowak, N. Burdzhiev, O. Petrov, E.S. Fischer, M. Etzkorn, V. Gelev, H. Arthanari, Local Deuteration Enables NMR Observation of Methyl Groups in Proteins from Eukaryotic and Cell-Free Expression Systems, Angew. Chem. Int. Ed. 60 (2021) 13783–13787. 10.1002/anie.202016070.

[48] J. Klopp, A. Winterhalter, R. Gébleux, D. Scherer-Becker, C. Ostermeier, A.D. Gossert, Cost-effective large-scale expression of proteins for NMR studies, J Biomol NMR. 71 (2018) 247–262. 10.1007/s10858-018-0179-0.

[49] E.L. Ulrich, H. Akutsu, J.F. Doreleijers, Y. Harano, Y.E. Ioannidis, J. Lin, M. Livny, S. Mading, D. Maziuk, Z. Miller, E. Nakatani, C.F. Schulte, D.E. Tolmie, R. Kent Wenger, H. Yao, J.L. Markley, BioMagResBank, Nucleic Acids Research. 36 (2007) D402–D408. 10.1093/nar/gkm957.

[50] P. Felig, The glucose-alanine cycle, Metabolism. 22 (1973) 179–207. 10.1016/0026-0495(73)90269-2.

[51] D.T. Wong, R.W. Fuller, B.B. Molloy, Inhibition of amino acid transaminases by L-cycloserine, Advances in Enzyme Regulation. 11 (1973) 139–154.

[52] P.W. Coote, S.A. Robson, A. Dubey, A. Boeszoermenyi, M. Zhao, G. Wagner, H. Arthanari, Optimal control theory enables homonuclear decoupling without Bloch– Siegert shifts in NMR spectroscopy, Nat Commun. 9 (2018) 3014. 10.1038/s41467-018-05400-4.

[53] T. Warne, M.J. Serrano-Vega, J.G. Baker, R. Moukhametzianov, P.C. Edwards, R. Henderson, A.G.W. Leslie, C.G. Tate, G.F.X. Schertler, Structure of a β1-adrenergic G-protein-coupled receptor, Nature. 454 (2008) 486–491. 10.1038/nature07101.

[54] E.R. Weiss, S. Osawa, W. Shi, C.D. Dickerson, Effects of Carboxy-Terminal Truncation on the Stability and G Protein-Coupling Activity of Bovine Rhodopsin, Biochemistry. 33 (1994) 7587–7593. 10.1021/bi00190a011.

[55] X. Deupi, P. Edwards, A. Singhal, B. Nickle, D. Oprian, G. Schertler, J. Standfuss, Stabilized G protein binding site in the structure of constitutively active metarhodopsin-II, Proc. Natl. Acad. Sci. U.S.A. 109 (2012) 119–124. 10.1073/pnas.1114089108.

[56] C.-J. Tsai, F. Pamula, R. Nehmé, J. Mühle, T. Weinert, T. Flock, P. Nogly, P.C. Edwards, B. Carpenter, T. Gruhl, P. Ma, X. Deupi, J. Standfuss, C.G. Tate, G.F.X. Schertler, Crystal structure of rhodopsin in complex with a mini-G o sheds light on the principles of G protein selectivity, Sci. Adv. 4 (2018) eaat7052. 10.1126/sciadv.aat7052.

[57] I. Shimada, T. Ueda, Y. Kofuku, M.T. Eddy, K. Wüthrich, GPCR drug discovery: integrating solution NMR data with crystal and cryo-EM structures, Nat Rev Drug Discov. 18 (2019) 59–82. 10.1038/nrd.2018.180.

[58] Y. Kofuku, T. Ueda, J. Okude, Y. Shiraishi, K. Kondo, M. Maeda, H. Tsujishita, I. Shimada, Efficacy of the β2-adrenergic receptor is determined by conformational equilibrium in the transmembrane region, Nature Communications. 3 (2012) 1045. 10.1038/ncomms2046.

[59] M.T. Eddy, M.-Y. Lee, Z.-G. Gao, K.L. White, T. Didenko, R. Horst, M. Audet, P. Stanczak, K.M. McClary, G.W. Han, K.A. Jacobson, R.C. Stevens, K. Wüthrich, Allosteric Coupling of Drug Binding and Intracellular Signaling in the A2A Adenosine Receptor, Cell. 172 (2018) 68–80.e12. 10.1016/j.cell.2017.12.004.

[60] S. Isogai, X. Deupi, C. Opitz, F.M. Heydenreich, C.-J. Tsai, F. Brueckner, G.F.X. Schertler, D.B. Veprintsev, S. Grzesiek, Backbone NMR reveals allosteric signal transduction networks in the β1-adrenergic receptor, Nature. 530 (2016) 237–241. 10.1038/nature16577.

[61] L. Wang, W.J. Zuercher, T.G. Consler, M.H. Lambert, A.B. Miller, L.A. Orband-Miller, D.D. McKee, T.M. Willson, R.T. Nolte, X-ray Crystal Structures of the Estrogen-related Receptor-γ Ligand Binding Domain in Three Functional States Reveal the Molecular Basis of Small Molecule Regulation, Journal of Biological Chemistry. 281 (2006) 37773– 37781. 10.1074/jbc.M608410200.

[62] F. Hagn, M. Etzkorn, T. Raschle, G. Wagner, Optimized Phospholipid Bilayer Nanodiscs Facilitate High-Resolution Structure Determination of Membrane Proteins, J. Am. Chem. Soc. 135 (2013) 1919–1925. 10.1021/ja310901f.

[63] T.A. Egorova-Zachernyuk, G.J.C.G.M. Bosman, A.M.A. Pistorius, W.J. DeGrip, Production of yeastolates for uniform stable isotope labelling in eukaryotic cell culture, Applied Microbiology and Biotechnology. 84 (2009) 575–581. 10.1007/s00253-009-2063-z.

