## Supplementary Information for "Human Cells for Human Proteins: Isotope Labeling in Mammalian Cells for Functional NMR Studies of Disease-Relevant Proteins"

### 1. Protocol for Uniform $^{15}$N-labelling in HEK293 Cells

This protocol is based on HEK293S GnTI$^-$ or HEK293T cells grown in suspension using V3 medium (V3-I, 500 ml, Bioconcept). Ensure that cells have high viability > 98% and are in logarithmic growth phase (cell density < $1.6 \cdot 10^6$ cells/ml for HEK293 in V3 medium) before starting with the protocol. In this example $^{15}$N-labeled yeast extract is used. Other amino acid extracts like algal extracts (Celtone, CIL or ISOGRO, Sigma) or in-house *E. coli* extracts can also be used but most likely will require prior delipidation.

All manipulations are carried out in sterile fashion under a biosafety cabinet.

**Day 1 evening: Transfer of cells into amino acid-poor medium and incubation for consumption of unlabelled amino acids**

1.1 Place V3 medium w/o yeastolate = V3$^{\ominus}$ (V3-702-I, Bioconcept) in a water bath set to 37°C

⇒ *Preheat medium to prevent slowed-down cell growth and clumping of suspension cells due to cold shock.*

1.2 Determine the cell count in the culture

1.3 Transfer $10^9$ cells from the cell culture into sterile centrifuge tubes

⇒ *Use sterile 250 mL Corning™ polypropylene large volume centrifuge tubes (Sigma CLS430776 or VWR 525-1597). Support cushions CLS430236 are needed for proper operation (Sigma CLS430236 or 3D printed https://www.thingiverse.com/thing:5162957) .*

⇒ *Calculate for a starting concentration of cell culture in new medium of $10^6$ cells $ml^{-1}$ at point 1.6 below.*

1.4 Centrifuge for 3 min at ≤ 125 g, 37°C

⇒ *Pre-warm the rotor to 37°C in an incubator before use*

1.5 Dispose supernatant by careful decanting

1.6 Re-dissolve pellet in 1 l of pre-warmed V3$^{\ominus}$ by gentle pipetting and transfer to a 5 l Erlenmeyer flask

⇒ *Use a 2 l flask for 0.5 l of culture to ensure sufficient gas exchange*

⇒ *As noted above, the target is a cell density of $10^6$ cells $ml^{-1}$.*

1.7 Place newly prepared culture in shaker-incubator (37°C, 5% $CO_2$, 100 rpm) and incubate for 16 h

⇒ *V3$^{\ominus}$ medium w/o yeastolate contains small amounts of amino acids, typically 10–50 mg of each amino acid. During this 16 h incubation period, the unlabeled amino acids are consumed. Do not let cells grow for longer than 18 h in this medium as it leads to starvation and lower final protein yields*

**Day 2 morning: Addition of labelled extract to cell culture.**

2.1 Remove 200 ml of the culture and return the rest of the culture to the shaker

2.2 Centrifuge the 200 ml of culture for 3 min at ≤ 125 g, 37°C

2.3 Decant the supernatant into a sterile flask

2.4 Re-dissolve the pellet in the remaining culture and return it to the incubator with shaker (37°C, 5% $CO_2$, 100 rpm)

2.5 Add 5 g $^{15}N$-labelled yeast extract per liter (CN700P, Cortecnet or 405304100, Silantes) and 10 mM $^{15}NH_4Cl$ (544.8 mg/L, Sigma) to the reserved supernatant and dissolve completely, sonicate if necessary

⇒ *In the acid hydrolysis performed in the production of amino acid extracts from simpler organisms (yeast, algae) glutamine is converted to glutamate, and asparagine to aspartate. The added $^{15}NH_4Cl$ is used by the cells to convert glutamate to labelled glutamine and resynthesize asparagine*

2.6 Sterilize the medium with added yeast extract by filtration (Steritop, Merck, 0.2 µm pore size)

2.7 Return the sterilized medium with added yeast extract and $^{15}NH_4Cl$ to the cell culture and incubate for approximately 2 h. (37 °C, 5% $CO_2$, 100 rpm)

⇒ *Wait 2 h to reduce stress to the cells before induction*

2.8 After 2 h, induce the cells by adding tetracycline (2 µg/ml) and sodium butyrate (5 mM)

**Day 4 or 5: Harvest**

4.1 Centrifuge culture for 10 min at ≤ 125 g, 4°C

4.2 Discard all but approx. 40 ml of the supernatant

4.3 Re-dissolve pellet and transfer to a 50 ml Falcon tube

⇒ *Increased yield by transferring cells in solution instead of scraping out the pellet*

4.4 Centrifuge for 10 min at ≤ 125 g, 4°C

4.5 Discard supernatant

4.6 Store pellet at –80°C until further use

#### 2. Protocol for Amino Acid specific labelling in HEK293 Cells

This protocol is based on HEK293S GnTI⁻ cells grown in suspension using V3 medium (V3-I, Bioconcept). Ensure that cells have high viability > 98% and are in logarithmic growth phase (cell density approx. $10^6$ cells/ml) before starting with the protocol.

All manipulations are carried out in sterile fashion under a sterile hood.

##### Day 1 evening: Transfer of cells into amino acid specific labelled medium

1.1 Prepare expression medium as described in Table 1, sterilize the medium by filtration using a 0.22 µm Steritop filtration unit (Merck), and pre-warm the medium to 37°C in a water bath

⇒ *Preheat medium to prevent slowed-down cell growth due to cold shock*

1.2 Determine the cell count in the culture

1.3 Transfer the required volume for the labelling culture from the current culture into sterile centrifuge tubes (starting concentration for cell culture in the labelling medium should be approximately $10^6$ cells $ml^{-1}$)

⇒ *Use sterile 250 mL Corning™ polypropylene large volume centrifuge tubes (Sigma CLS430776) or alternative from VWR (Cat. No. 525-1597). Support cushions CLS430236 are needed for proper operation.*

1.4 Centrifuge for 5 min at 100 g, 37°C

⇒ *Pre-warm the rotor to 37°C in an incubator before use*

1.5 Dispose supernatant by careful decanting

1.6 Re-dissolve pellet in pre-heated labeling medium by gentle pipetting and transfer suspension and remaining medium to a 5 l Erlenmeyer flask.

⇒ *Use a 5 l flask for 1 l of culture, 2 l flask for 0.5 l of culture*

1.7 Place newly prepared culture in shaker-incubator (37°C, 5% $CO_2$, 100 rpm)

⇒ *Let cells grow for 2.5 days*

##### Day 4 or 5: Harvest

4.1 Centrifuge culture for 10 min at ≤ 125 g, 4°C

4.2 Discard all but 50 ml of the supernatant

4.3 Re-dissolve pellet and transfer to a 50 ml Falcon tube

⇒ *Increased yield by transferring cells in solution instead of scraping out the pellet*

4.4 Centrifuge for 10 min at ≤ 125 g, 4°C

4.5 Discard supernatant

4.6 Store pellet at –80°C until further use

Table 1: Recipe for the preparation of media for the labelling of the methyl group bearing amino acids alanine, isoleucine, leucine, valine and for phenylalanine. All quantities are given for the preparation of 1 l of medium.

| **Component** | **Ala** | **Ile** | **Leu** | **Val** | **Phe** |
|---|---:|---:|---:|---:|---:|
| *Amino Acid Mix* | | | | | |
| L-Alanine | | 95 mg | 95 mg | 95 mg | 95 mg |
| L-Arginine | 60 mg | 60 mg | 60 mg | 60 mg | 60 mg |
| L-Asparagine | 110 mg | 110 mg | 110 mg | 110 mg | 110 mg |
| L-Cysteine | 15 mg | 15 mg | 15 mg | 15 mg | 15 mg |
| L-Glutamate | 155 mg | 155 mg | 155 mg | 155 mg | 155 mg |
| Glycine | 55 mg | 55 mg | 55 mg | 55 mg | 55 mg |
| L-Histidine | 30 mg | 30 mg | 30 mg | 30 mg | 30 mg |
| L-Isoleucine | 60 mg | | 60 mg | 60 mg | 60 mg |
| L-Leucine | 115 mg | 115 mg | | 450* mg | 115mg |
| L-Lysine | 80 mg | 80 mg | 80 mg | 80 mg | 80 mg |
| L-Methionine | 25 mg | 25 mg | 25 mg | 25 mg | 25 mg |
| L-Phenylalanine | 60 mg | 60 mg | 60 mg | 60 mg | |
| L-Proline | 60 mg | 60 mg | 60 mg | 60 mg | 60 mg |
| L-Serine | 50 mg | 50 mg | 50 mg | 50 mg | 50 mg |
| L-Threonine | 55 mg | 55 mg | 55 mg | 55 mg | 55 mg |
| L-Tryptophan | 15 mg | 15 mg | 15 mg | 15 mg | 15 mg |
| L-Tyrosine | 50 mg | 50 mg | 50 mg | 50 mg | 50 mg |
| L-Valine | 75 mg | 75 mg | 75 mg | | 75 mg |
| Yeast Extract | 500 mg | 500 mg | 500 mg | 500 mg | 500 mg |
| *Labelled Amino Acids and Inhibitors* | | | | | |
| $^{13}C^{\beta}$-Alanine | 250 mg | | | | |
| L-Cycloserine | 5 mg | | | | |
| $^{13}C_6$-Isoleucine | | 250 mg | | | |
| $^{13}C^{\delta}$-α-keto-isocaproate | | | 160 mg | | |
| *or* $^{13}C^{\delta}$-Leucine | | | *or* 230 mg | | |
| $^{13}C^{\gamma}$,$^{2}H$ α-keto-isovalerate | | | | 160* mg | |
| *or* $^{13}C^{\gamma}$-Valine | | | | *or* 150 mg | |
| $^{13}C^{\gamma}$-Phenylalanine | | | | | 160 mg |
| *Base Medium and Further Components* | | | | | |
| Fetal Calf Serum (FCS/FBS) | | | 50 ml | | |
| Tetracycline 2 mg/ml | | | 1 ml | | |
| Sodium Butyrate 500 mM | | | 10 ml | | |
| V3 Medium w/o Yeastolate | | | 939 ml | | |

* When using the precursor α-keto-isovalerate, scrambling to leucine occurs. This can be prevented by adding an excess of unlabeled leucine to the medium, i.e. 450 mg instead of 115 mg/l.

#### 3. Protocol for preparation of labelled amino acid extracts from *E. coli*

For this approach, *E. coli* cells with the desired labeling pattern are grown, and subsequently an amino acid extract is prepared for feeding mammalian cells. Typically, *E. coli* cells are already used for expressing a target protein, which is purified, and only the discarded lysate and flow-through fractions from the purification are used for preparation of the isotope labeled amino acid extract, thus economizing on isotope labeled compounds. ILV-methyl labeling will be used as an example in this protocol. In this case, cells however must be grown from the start in medium containing isotope labeled precursors (α-keto-isovalerate and α-keto-butyrate, at 180 and 160 mg/l, respectively). If precursors are added only at the timepoint of induction of the *E. coli* culture, the resulting overall incorporation is too low (~60–70%).

##### Day 1 : Lysis and first digestion

1.1 Pool lysate pellets and flow through from purification from uniform $^{15}N$, Leuδ-[$^{13}CH_3$], Valγ-[$^{13}CH_3$] and Ileδ$_1$-[$^{13}CH_3$] labelled *E. coli* cells (ca. 150 ml from 1 l of culture)

1.2 Heat solution to 92°C for 25 min and centrifuge at 5000 rpm for 15 min at room temperature (Beckmann, JLA 8.1000 rotor).

1.3 Discard the supernatant and resuspend the pellet in 50 ml water. Adjust the pH to 7 (no buffer is added, as the biological material acts as a buffer itself).

1.4 Add 6 mg papain (from *Carica papaya*, Fluka 76220) and let the suspension be digested overnight at room temperature and under constant stirring.

##### Day 2 : Further digestions

2.1 Run a second, high-temperature digestion by adding 6 mg of papain and incubate for 3 hours at 65°C and 600 rpm on a thermal shaker (Thermomixer comfort, Eppendorf).

2.2 For a third digestion, add 15 mg of bromelain (from pineapple stem, Sigma-Aldrich B4882) adjust the pH to 5.5, and incubate at 50°C and 600 rpm for 4 hours.

2.3 For the fourth digestion, adjust the pH to 1–1.5 and supplement the solution with 12 mg of pepsin (from hog stomach, Fluka 77163). Incubate at 37°C overnight.

##### Day 3 : Purification of extract

3.1 Heat the sample to 92°C for 20 minutes, cool down to room temperature and filter through an MN615 1/4 filter paper and a 0.45 µm membrane filter (Millipore HAWP04700)

3.2 Add a 3-fold excess of chloroform and vigorously stir the biphasic suspension on a magnetic stirrer overnight.

##### Day 4 : Extraction of amino acids and lyophilization

4.1 Let the phases separate and discard the lower organic phase using a separation funnel. Take special care not to collect any of the foam-like border phases.

4.2 Finally, lyophilize the aqueous phase to obtain a dry amino acid extract (~1 g per liter of original *E. coli* culture)

#### 4. Protocol for delipidation of cell extracts

This protocol is based on the modified Bligh and Dyer method for lipid extraction [1].

##### Day 1 : Dissolution of cell extract

1.1 Dissolve extract (e.g. ISOGRO-$^{13}C$,$^{15}N$ Powder-Growth Medium, Sigma Aldrich) in the smallest possible amount of water (dd$H_2O$).

1.2 Add chloroform and methanol to the solution in a 1 : 1.25 : 2.5 v/v ratio, where 1 corresponds to the volume of water.

1.3 Stir resulting monophasic solution overnight at room temperature.

##### Day 2 : Separation of organic and aqueous phases

2.1 Add water and chloroform to achieve a final v/v ratio of water, methanol and chloroform of 2.25 : 2.5 : 2.5. Two distinct phases should now be visible. Let the system separate for >30 min.

2.2 Separate the lower organic phase using a separation funnel. This phase can be discarded.

2.3 Reduce the volume of the aqueous phase in a rotary evaporator to less than 50 ml.

2.4 Freeze and lyophilize the concentrate to obtain water- and methanol-free dry powder.
